# ImaGene: A web-based software platform for tumor radiogenomic evaluation and reporting

**DOI:** 10.1101/2021.12.02.470994

**Authors:** Shrey S. Sukhadia, Aayush Tyagi, Vivek Venkataraman, Pritam Mukherjee, AP Prathosh, Mayur D. Divate, Olivier Gevaert, Shivashankar H. Nagaraj

## Abstract

The field of radiomics has undergone several advancements in approaches to uncovering hidden quantitative features from tumor imaging data for use in guiding clinical decision-making for cancer patients. Radiographic imaging techniques provide insight into the imaging features of tumor regions of interest (ROIs), while immunohistochemistry and sequencing techniques performed on biopsy samples yield omics data. Potential associations between tumor genotype and phenotype can be identified from imaging and omics data via traditional correlation analysis, as well as through artificial intelligence (AI) models. However, at present the radiogenomics community lacks a unified software platform for which to conduct such analyses in a reproducible manner.

To address this gap, we propose ImaGene, a web-based platform that takes tumor omics and imaging data sets as input, performs correlation analysis between them, and constructs AI models (optionally using only those features found to exhibit statistically significant correlation with some element of the opposing dataset). ImaGene has several modifiable configuration parameters, providing users complete control over their analysis. For each run, ImaGene produces a comprehensive report displaying a number of intuitive model diagnostics.

To demonstrate the utility of ImaGen**e,** exploratory studies surrounding Invasive Breast Carcinoma (IBC) and Head and Neck Squamous Cell Carcinoma (HNSCC) on datasets acquired from public databases are conducted. Potential associations are identified between several imaging features and 6 genes: CRABP1, SMTNL2, FABP1, HAS1, FAM163A and DSG1 for IBC, and 4 genes: CEACAM6, NANOG, ACSM2B, and UPK2 for HNSCC.

In summary, the software provides researchers with a transparent tool for which to begin radiogenomic analysis and explore possible further directions in their research. We anticipate that ImaGen**e** will become the standard platform for tumor analyses in the field of radiogenomics due to its ease of use, flexibility, and reproducibility, and that it can serve as an enabling centrepoint for an emerging radiogenomic knowledge base.

## INTRODUCTION

Diagnostic imaging techniques are routinely used in clinics and laboratories for the identification of tumor severity and progression (Bodalal et al. 2019). The most common such techniques include computed tomography (CT), magnetic resonance imaging (MRI), and positron emission tomography (PET). These techniques yield high-quality digital images for tumor assessment that are also suitable for building databases that curate patient data for the purposes of research reproducibility and reuse (Diaz et al. 2021; Freymann et al. 2012; Gillies et al. 2016; Prior et al. 2017).

When used to assess tumors, diagnostic imaging techniques produce images that are examined by radiologists to segment (or select) tumor regions of interest (ROIs) that are believed to represent core sections of tumor on images. Measurements such as spatial, volumetric, textural and intensity-based are extracted from image-based ROIs, and analyzed using several techniques in correlation-analysis and artificial intelligence (AI) (Pfaehler et al. 2019; van Griethuysen et al. 2017). Consequently, the best portion of ROI is marked on the image and the corresponding tissue material is biopsied from patient’s body and subjected to histopathological examination (Lee et al., 2013; Rice et al., 2017) and omics-based assessment (González-Reymúndez and Vázquez 2020; Y. Liu et al. 2018; Nasrallah et al. 2019; Peng et al. 2015; W. Wang et al. 2018). Omics-based investigations yield information such as gene expression, copy number variations (CNVs) or other structural variants (SVs), single nucleotide variations (SNVs), and DNA methylation scores for genes that regulate biological processes within the biopsied tissue material (Anker and Chae 2015; Balassiano et al. 2011; Chen et al. 2013; Park et al. 2010).

Despite the widespread use of tumor biopsies as a means of establishing important diagnostic information pertaining to specific ROIs, several studies have reported an underestimation of adverse pathologies in up to 23% of the samples due to spatial sampling bias (Aihara et al. 1994; Kvale et al. 2009; Martin-Gonzalez et al. 2020; Siddiqui et al. 2015; Smith et al. 2019). This may be attributed to the spatial heterogeneity of morphological growth patterns (Aihara et al. 1994; Bosaily et al. 2016; Boutros et al. 2015) as well as genetic heterogeneity within the cancerous lesions and may result in the underestimation of tumor severity, leading to an increase in the potential for tumor-related adverse events in these patients (Aihara et al. 1994; Incoronato et al. 2020; Martin-Gonzalez et al. 2020; Sottoriva et al. 2013). Therefore, combining imaging information of tumors with their omics profile could explain the tumor heterogeneity better than just with imaging alone. One such method would be to correlate data from image-based tests and omics-based investigations in order to improve the quality of diagnosis (Bakr et al. 2018; Gevaert et al. 2012; Gevaert et al. 2014; Lo Gullo et al. 2020; Martin-Gonzalez et al. 2020; Zhou et al. 2018) and to provide more reliable tumor ROIs that can be used for tissue sampling when performing biopsies (Gillies et al. 2016; Martin-Gonzalez et al. 2020). Further, AI models built with flexibility in performing prior filtering of features using statistical correlation analysis may allude crucial associations that may lead to improved tissue biopsies in patients (Ashraf et al. 2014; Bodalal et al. 2019; Chitalia et al. 2020). Such models may also reduce or obviate the need for tissue biopsy for tumor-assessment in near future (Bodalal et al. 2019).

The field of imaging genomics or radiogenomics focuses on finding associations between radiomic characteristics of tissue ROIs and molecular characteristics such as genomic, transcriptomic, and proteomic profiles of tumor cells. The Cancer Imaging Archive (TCIA) recently conducted a locoregional study using MRI images of glioblastoma tissues by generating heat maps corresponding to both core and boundary regions of specific tumor ROIs, revealing substantial genetic heterogeneity (Depeursinge et al. 2018). The gene expression profiles of each of the sampled regions were correlated with known information regarding the genes involved in multiple pathways potentially leading to oncogenesis (Depeursinge et al. 2018). Another recent study investigated gene expression profiles of pancreatic ductal adenocarcinoma (PDAC) ROIs using CT images by comparing the radiomic features of these ROIs with their genotypes and stromal content (Attiyeh et al. 2019). This investigation yielded a radiogenomic model that provided important information regarding the number of altered genes in the ROIs and SMAD4 gene expression status in association with radiomic features. Stromal content was also correlated with radiomic and genomic features of the ROIs, thereby indicating that stromal information can guide decision-making for PDAC samples (Attiyeh et al. 2019).

Radiogenomic investigations typically use a combination of machine learning and statistical methods to detect correlations between the radiomic and genomic features of tumor regions. These methods are unique for each study and most studies do not provide the details of the codes and parameters used for these investigations, thereby limiting their reproducibility (Brito et al., 2020; Colen et al., 2014). Additionally, they do not offer users with enough algorithmic flexibility, for example, using AI modelling with or without prior correlations filtering of the features, the approaches practiced commonly in the field so far (Ashraf et al. 2014; Chitalia et al. 2020; Trivizakis et al. 2020). There is currently a need for the development of a sophisticated, user-friendly, web-based software platform that can perform both correlation analysis and modeling operations using AI methods and provide trained models that could be used for rigorous training and testing associations of imaging features with the omics profiles of tumor ROIs (Bodalal et al. 2019; Colen et al. 2014). There is also a need for a transparent algorithm, the parameters of which can be substantially exposed and altered in order to obtain comprehensive and self-explanatory reports that can be used for radiogenomics research and hopefully cancer diagnosis post sufficient validations through an unified software platform.

At present, there are several software tools available that can detect associations between imaging characteristics and gene regulatory networks in tissue samples, including Imaging-Amaretto and Imaging-Community Amaretto. However, these tools neither have a sophisticated, user-friendly, web-based platform that can allow for the straightforward manipulation of experimental parameters, nor do they provide comprehensive reports that describe statistical correlations, their performance metrics (such as correlation co-efficients, intuitive plots such as clustered heat maps, and the AI based prediction and/or classification of labelled data along with the metrices such as RMSE:STDEV ratio, R-square and area under the receiver operating curve (AUROC / AUC) that explain the performance models in predicting and/or classifying either omics or imaging data from either imaging or omics data, respectively (Gevaert et al. 2020). Additionally, users require a flexibility to select from a variety of AI model types (mainly regression based used in radigenomic domain so far) for training and testing, and consequently post analyzing or comparing results using quality control metrices such as RMSE:Stdev, R-square and AUC to arrive at reliable imaging-omics associations (Gevaert et al. 2020). Multi-omics Statistical Approaches (MuSA) is another tool used for radiogenomic studies that provides correlation heat maps and principal component analysis (PCA) plots. However, it lacks a feature prediction ability imparted by AI-based methods (Zanfardino et al. 2021). All of these tools also fail to adhere to the Findability, Accessibility, Interoperability, and Reusability (FAIR) principles that allow users to store, track, and analyse their data through both individual steps and the entire experiment (Wilkinson et al. 2016).

In order to address this need for a universal platform that can conduct correlation analysis between imaging and omics-based features of tumor ROIs and build AI models based on or off of such correlated features, we developed ImaGene, which is a software platform that integrates statistical and AI techniques to facilitate tumor radiogenomic analyses. ImaGene is a web-based platform that facilitates systematic radiogenomic analysis using various statistical and AI parameters that allow users to configure their experiments and perform appropriate iterations thereof. The end result of this platform is an HTML document that describes how the experiment was executed, the parameters that were used, and the resulting associations in the form of correlation plots and performance metrics (and plots) of AI models such as Mean Squared Error (MSE) and Root Mean Square Error: Standard deviation (RMSE:Stdev) ratio. The AI piece consists of regression models such as linear, regularized regression (LASSO and elastic net along with their respective multitask versions), and decision trees. It further conducts classification of predicted labels at various decision thresholds yielding AUC values at such thresholds. Through these outputs, ImaGene allows users to perform their radiogenomic experiments more intuitively and helps them establish unambiguous conclusions that can highlight data-driven directions for future research.

### Features of ImaGene

ImaGene has been deployed as a Graphical User Interface (GUI) on the Amazon Web Service (AWS) servers of Queensland University of Technology (QUT), and is also available as an open-access website ‘www.ImaGene.pgxguide.org’ (Figure 3). Users can log in to the website using common credentials (Table 2) or register for a free account. Alternatively, users can download ImaGene from GitHub (https://github.com/skr1/Imagene.git) and operate it on Linux Operating System (OS) using a Command-Line Interface (CLI). Once downloaded, users can set various configuration parameters available on the platform and run their experiments (Figure 3). ImaGene comprises of four modules: a) data pre-processing, b) correlation Analysis, c) machine learning (ML), and d) reporting (Figure 1). The code for the software has been written in Python, and it utilizes several libraries such as scikit-learn (Pedregosa et al. 2011), matplotlib, seaborn and importr along with custom functions written to follow a systematic approach to analyze and pin point meaningful associations between imaging and omics features.

**Figure 1.**
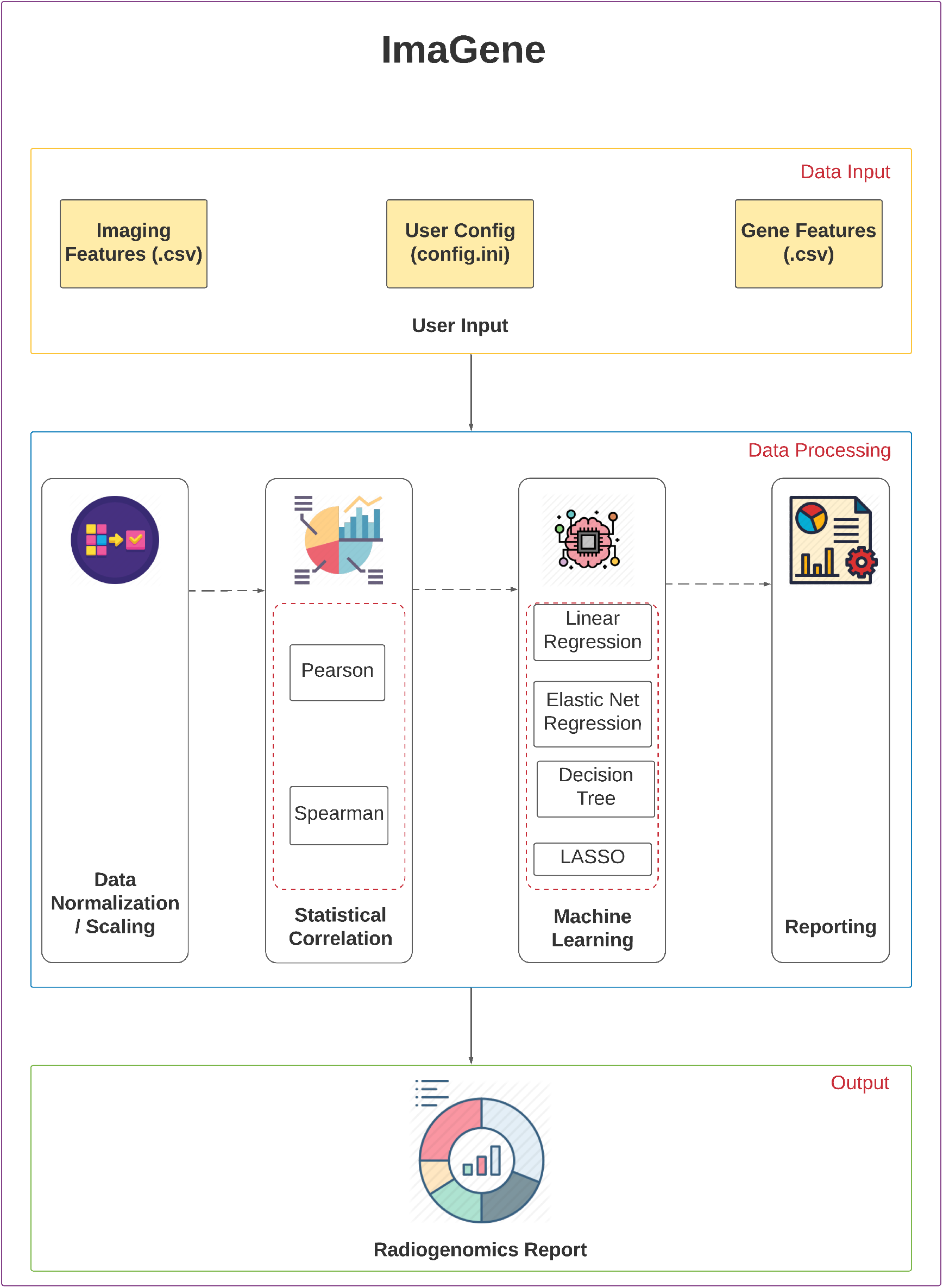
ImaGene’s Schema depicting all its components.

**Figure 2.**
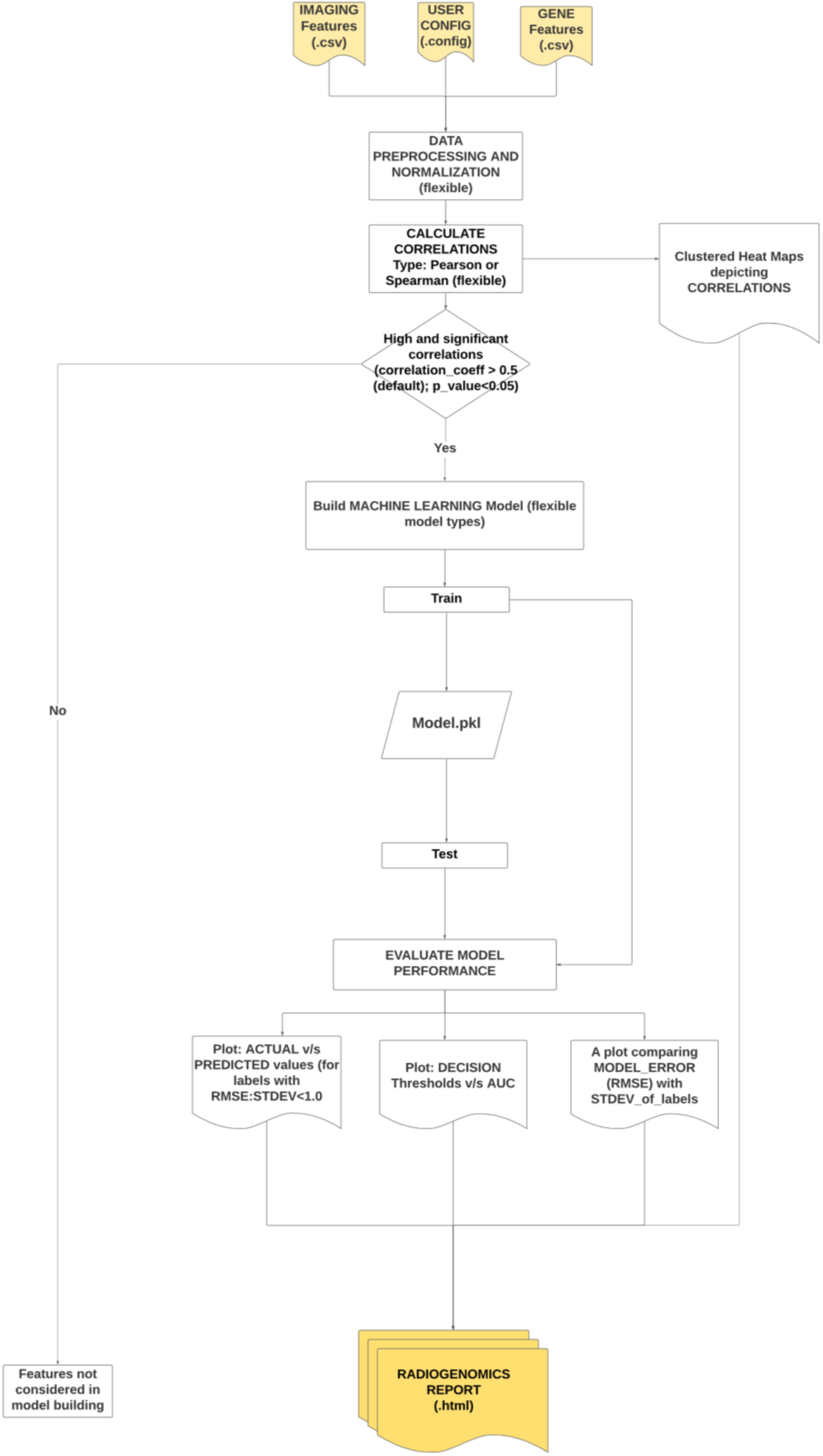
Calculating Correlations and conducting Training and Testing of Machine Learning Models.

**Figure 3:**
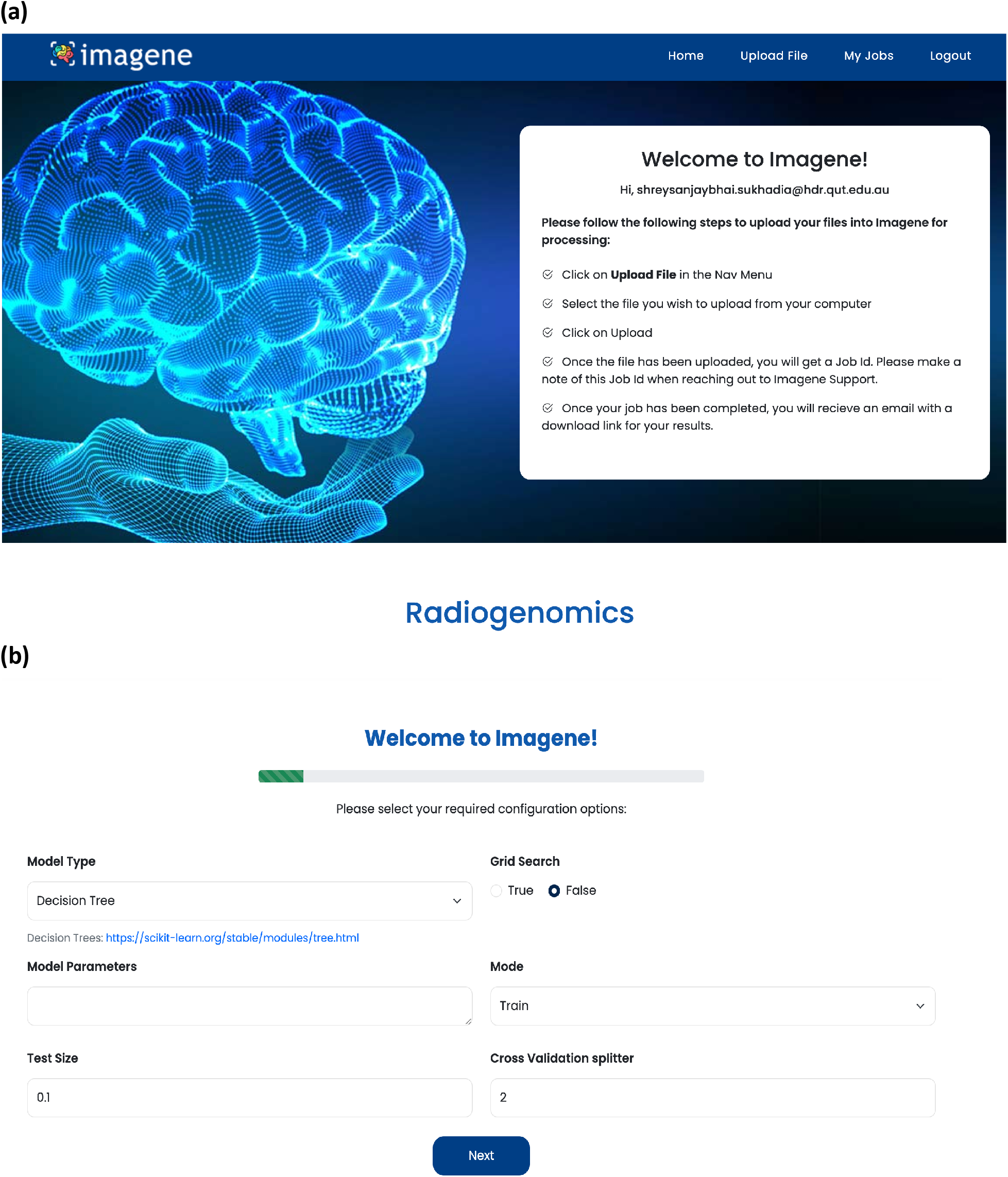
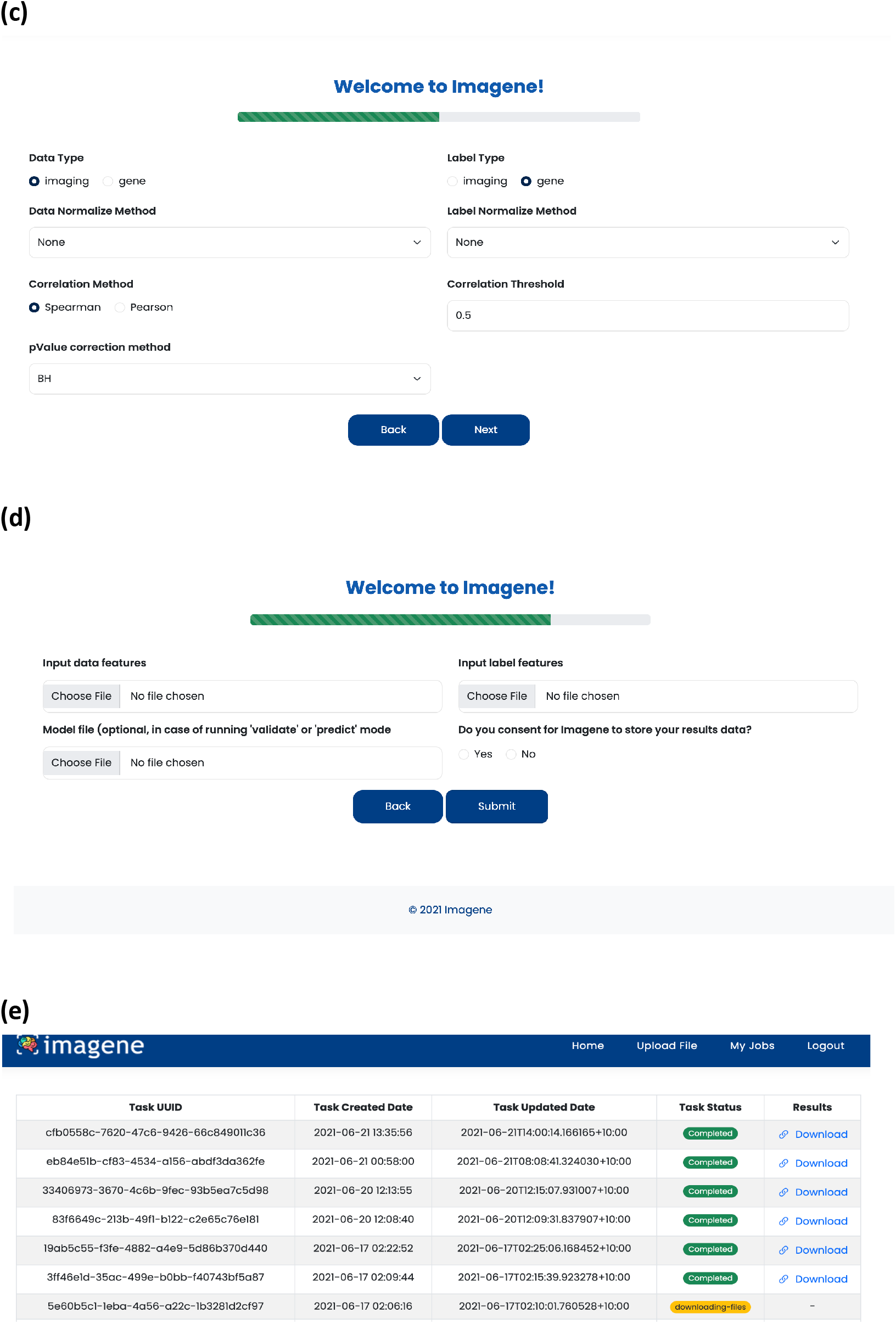
a) Welcome page for ImaGene, b-c) Parameter selection for ImaGene d) Selecting radiomic and genomic files, and model file depending on the mode of operation e) Job tracking and results download page.

**Table 1:** Configuration Parameters for ImaGene

| Configuration Parameters | Values |
| --- | --- |
| Correlation Method | Default: Spearman (options: Pearson) |
| Correlation Threshold | Default: 0.5 (flexible) |
| p- value correction method | Default:BH (options: holm, hochberg, hommel, bonferroni, BH, BY, fdr) |
| Data feature-type | Default:Imaging (option: Gene) |
| Label feature-type | Gene (option: Imaging) |
| Train size | 0.9 (flexible) |
| Test size | 0.1 (flexible) |
| Normalization method | Default: Stand_scaler (options: min_max , Stand_scaler , zscore, MaxAbsScaler) |
| Mode | Default: Train (options: Train, validate, predict) |
| Model Type | Default: Decision Tree (options: Linear Regression, Linear Model aka Elastic-Net, LASSO, Multi task LASSO and Multi task Elastic-Net) |
| Grid Search | default: False (options: True, False) |
| Cross-Validation splitter | cv: 2 (flexible) |
| <Section for other model parameters for each model type> | Could be filled with as many model parameters as possible or kept empty. |
| <Section for Grid Search parameters for each model type> | Could be filled with as many grid search parameters as possible or kept empty. |

**Table2:**
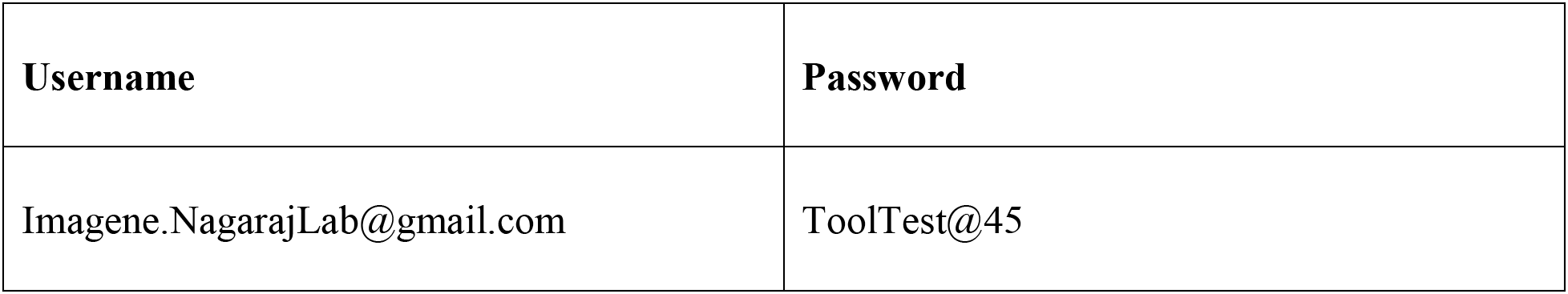
Common user credentials for ImaGene web-platform

For analytical operations, ImaGene requires two input files, each in Comma-Separated Value (CSV) format – one containing imaging features (and their measurements) and another enhousing omics features (and their measurements) for a set of tumor samples. The imaging features can be acquired from feature extraction software such as PyRadiomics, LIFEx, or RaCaT by processing the tumor images using the respective segmentation labels (Koçak et al. 2019; Pfaehler et al. 2019; van Griethuysen et al. 2017). The omics features can be acquired from studies conducted on tumor ROIs in biopsy samples processed in pathological laboratories, and may consist of data pertaining to gene expression, SV (including CNV), SNV, or DNA methylation scores.

Once the imaging and omics feature files are uploaded in the software, the names and order of samples therein are matched. Following this, the feature files are analyzed for different configuration parameters based on values specified by the user in the data pre-processing module (Table 1; Figure 1). The imaging and omics features undergo normalization or scaling based on the normalization method set by the user such as Standard Scaler, Max Absolute Scaler, MinMax Scaler, or Z Score Normalization (Table 1).

The data is then transferred to the correlation analysis module where either Pearson or Spearman test is conducted for imaging and omics features per user’s selection of the correlation method (Table 1). The outcome of the correlation test is a subset of imaging and omics features that are highly correlated with one another (correlation coefficient > 0.5). The correlation threshold configuration parameter allows users to specify this minimum correlation coefficient threshold (Table 1). For strong correlations found in the imaging and omics feature files, the default Benjamin-Hochberg p-value correction method (Table 1) is used to measure and adjust the p-values for false discovery rates. Features having significant correlations are presented as hierarchically clustered heat maps depicting three types of relationships between imaging and omics features – a) two-way univariate, b) univariate to multivariate, and c) two-way multivariate. These features (p_adjust < 0.05) are then transferred to the ML module for further processing.

In the ML module, the statistically correlated features are used to construct a definitive ML model based on the model type specified in the parameter settings (Table 1). Currently, the model types available in the platform are linear, regularized regression (LASSO and elastic net along with their respective multi-task versions), and decision trees. Users may specify their preferred model parameters (Table 1) depending on the options available in the Scikit-learn library (Pedregosa et al. 2011). Alternatively, if users are unsure of these parameters, they may leave this section empty or specify only a subset of parameters, in which case, default values for unspecified parameters will be used (Figure 3b). Users also have the flexibility to specify omics and imaging features as either data, which is the independent variable set (X) where, {*X* = {*x*_1_, *x*_2_… *x_n_*} or label, which is the dependent variable set (*Y*) where {*Y* = {*y*_1_, *y*_2_… *y_n_*} and which is predicted based on the independent variable set (*X*). ImaGene also provides users with an option to specify grid parameters for their experiments based on the options available in the Scikit-learn library, which can be achieved by setting the ‘grid search’ parameter to ‘True’ (Figure 3b), and by either mentioning the list of grid search parameters in the ‘Model Parameters’ section (Figure 3b) or by using the default options. Specifying grid search parameters allows users to train models using different settings for model parameters, finalizing a model with the best performing parameter settings.

The ML module uses either user-defined settings or default settings to construct an AI model that is trained using a training dataset. Following this, a K-fold cross-validation is performed to reduce data overfitting based on the value specified in the ‘Cross-Validation Splitter’ parameter (cv, default value = 2) (Table 1). Users can also specify the fraction of the total number of test samples that should be used for testing the model by entering their desired value in the ‘Test Size’ parameter (default value = 0.1) (Table 1, Figure 3b). Based on this setting, the test dataset is used for testing the model.

The test dataset is scored by calculating the negative mean square error (MSE) and the root mean square error (RMSE). RMSE is calculated for each label (*y*) in the dataset, and the ratio of RMSE “RMSE(*y*)” and standard deviation of the originally observed values “STDEV(*y*)” (RMSE: STDEV) represents the prediction error of the model for each label as a function of the standard error (standard deviation) in the distribution of the observed label values (y_observed_). Further, users could utilize the identity (1) below depicting the relationship between R-square (*R*^2^) and RMSE:Stdev ratio (*v*) to compute the *R*^2^. The *R*^2^ along with *v* provides users with two metrics that aid in assessing the reliability of various regression models in the context of residual variance for *y*_predict_ (i.e. SSL) and the total variance that exists in *y*_observed_ (i.e., SST).

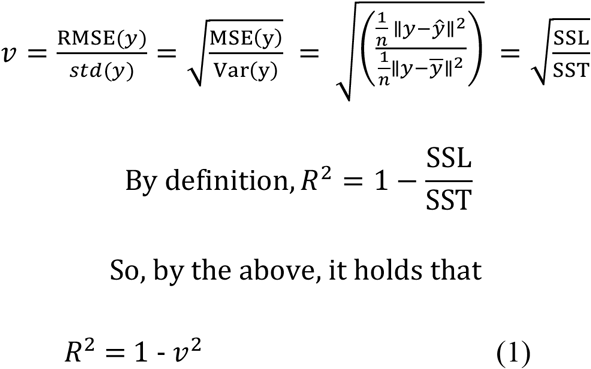

In order to evaluate the performance of a regression model when used to classify omics labels (such as genes) into groups of their high and low measures (for instance, high and low normalized gene-expressions which are in continuous form), such measures are first binarized for both y_observed_ and y_predict_ at various decision thresholds (aka normalized gene-expression cut-offs) for each and every gene label (*y*). Thus, if a normalized expression value belongs to a range of [0,1] increasing in steps of 0.1 (i.e. *dt* = Error! Bookmark not defined.), and if the value of the decision threshold is set to 0.5 (as an example), all the expression values falling below 0.5 will be set to 0 and the ones landing above 0.5 will be set to 1, for both y_observed_ and y_predict_ for every gene label (*y*). Such a binarization allows ImaGene to calculate AUCs using binarized y_observed_ and y_predict_ values for each label (gene) at various measurement (aka normalized gene-expression) cut-offs, which aids users evaluate the model’s ability to classify labels (genes) into groups of high and low measures (at definite thresholds) indicating them the measures at which the labels (genes) are more reliable to be predicted using imaging features as the dataset (Depeursinge et al. 2018). A plot of the AUC values versus decision thresholds is generated for all gene labels providing users with a visualized overview (along with the respective supplementary data files) indicative of a regressionmodel’s classification performance. This is very beneficial from biological perspective as users can determine expression cut-offs for a gene at which a regression-model in radiogenomics could be relied upon for predicting a gene into high or low expressed category using imaging features from a tumor ROI on the radiographic image.

Validation of the model can be conducted using multiple test datasets in validation mode (Supplementary Figure 2). The prediction mode of the platform can be used to perform omics predictions on new imaging datasets (Supplementary Figure 3). The ‘job tracking’ feature of ImaGene allows users to track the status of their experiments, and to download the reports and the supporting result files (along with intermediate calculated tables) of completed experiments using the provided download links (Figure 3e) on the ‘jobs’ page on the web platform.

### A Radiogenomics Report

Once an experimental run is completed, users can use the job tracking page to download a detailed HTML report. This report represents the alterations made to the imaging and omics data in each module of ImaGene in the form of scores and plots which help users acquire an overview of the process and deduce meaningful outcomes from the features data (Supplementary reports 1 and 2). The first section of the report provides details regarding the type of input data (i.e. imaging and omics features files), the number of sample entries common to both files, and the type of normalization method used for the input data. The second section provides hierarchically clustered heat maps depicting statistical correlations between the imaging and omics features. Features having a correlation coefficient greater than the default threshold of 0.5 and an FDR-adjusted p-value greater than the cutoff value of 0.05 are used to train user-defined AI models and set hyperparameters including the K-fold cross validation splitter (i.e. the ‘cv’ parameter). The next section describes the imaging and omics features that exhibited significant correlations and that were used to train the model along with the parameters that were used for training and testing the model. The next section is the ‘model interpretation’ section which provides metrics such as MSE, RMSE, and K-fold cross-validation scores of the model, and various plots such as a bar plot representing the RMSE: STDEV ratio for each label feature, a scatter plot representing true versus predicted values of labels having RMSE: STDEV ≤ 1, and a two-dimensional plot representing decision threshold values on the x-axis and AUC values on the y-axis to indicate the performance of the model in classifying labels (genes) into categories (high or low expression) at different cutoff values (normalized gene expressions – FPKMs) from 0 to 1. These metrices aid users determine the model’s reliability in predicting and/or classifying labels.

To demonstrate the performance and capabilities of ImaGene, we next conducted two case study analyses of tumor imaging data from The Cancer Imaging Archive (TCIA) and corresponding gene expression data from TCGA pertaining to two major types of cancer - Invasive Breast Carcinoma (IBC), and Head and Neck Squamous Cell Carcinoma (HNSCC). To clarify further, these use cases do not necessarily obligate the user to use ImaGene’s parameters similar to ours, and that users have flexibility to operate on same datasets using the parameters of their choice. The imaging (or radiomic) and gene expression datasets are provided as supplementary tables for IBC and HNSCC (i.e., Supplementary_table_BC-Gene_FPKM.csv, Supplementary_table_BC-Radiomic_features.csv, Supplementary_table_HNSCC-Gene_FPKM.csv andSupplementary_table_HNSCC-Radiomic_features.csv).

### Case Study 1: Invasive Breast Carcinoma (IBC)

The TCGA Breast Phenotype Research Group datasets available on the TCIA platform were accessed to extract 36 imaging features from tumor ROIs in MRI scans of 89 IBC patients using the TCIA NBIA Data Retriever software (Burnside et al. 2016; Clark et al. 2013; Li et al. 2016b; Li et al. 2016a; Zhu et al. 2015). A gene expression dataset comprised of FPKM (Fragments Per Kilobase of transcript per Million mapped reads) data was downloaded from the main TCGA-BRCA sample set of which the 89 patient samples formed a subset. FPKM data was screened for genes that had an FPKM value greater than or equal to 5 for at least 30 patients in the TCGA-BRCA cohort. In total, 976 genes were obtained that met with these criteria. This type of screening ensures reliable correlations between imaging and omics data (gene-FPKM data in this case study) such that these results can be leveraged for the generation of high-quality AI models capable or generating predictions or classifying gene expression levels based upon tumor ROI imaging features.

Once the initial screening of the downloaded dataset was complete, the imaging and gene-FPKM features were processed based upon user-defined ImaGene parameter settings. For example, the absolute Pearson-based correlation coefficient threshold was set to 0.7 for linear regression modeling downstream and 0.6 for other model types depending on number of features required for the satisfactory training of different types of AI models, the P-value correction method was set to Benjamin-Hochberg (BH), and the corrected p-value threshold was set to 0.05. Furthermore, the imaging features were set as data and gene-FPKM values were set as labels. For both data and labels, the normalization method was set to StandScaler, the test size was set to 0.2 (i.e. 20% of the dataset), and the K-fold cross-validation splitter (cv) was set to 3.

At the end of the experimental run using the abovementioned parameters, a comprehensive radiogenomics report was obtained for each model type that reported Linear Regression (LR), Decision Tree (DT), Multi-Task Lasso (MTL), and Multi-Task Elastic Net (MTEN) models (Supplementary Reports BC 1-6) that yielded the best results for the test dataset. The last two models were executed with two scenarios, i.e. with and without prior correlation analysis. These reports represented four normalized radiomics features – a) Signal Enhancement Ratio (SER) (K7), b) Size of the lesion/volume (S1), c) Surface area (S3), and d) Volume of the most enhancing voxels (S4) – that were first significantly correlated with the normalized FPKM values of a total of 25 genes as depicted in the reports (Supplementary Reports BC 1-4). Secondly, when the correlation coefficient threshold was relaxed to 0.6, an additional radiomics feature (Washout rate (K4)) and 5 additional genes appeared on the list of features exhibiting significant correlations (Supplementary Reports BC 2-4). Using this threshold and the expanded feature set, the training of the Decision Tree model was found to be enhanced. Lastly, the number of genes got reduced to more significant ones (only 11 out of 30) after application of several AI models to predict gene expressions from the above mentioned radiomic features.

The LR and DT models yielded test MSE values of 1.27 and 3.2 respectively, and respective K-fold cross-validation scores of 1.59 and 0.97 (Supplementary reports BC 1-2) during their training indicating low overfitting of the data by them. For both these models, ImaGene measured the AUC value for each gene label across the predicted absolute normalized FPKM decision threshold in the range of [0,1] (Table 3a–b, Supplementary Table BC LR2, and Supplementary Table BC DT4). This analysis revealed the CRABP1 and SMTNL2 genes to be strongly associated with the radiomics features with AUC in the range of [0.97, 1.0] at multiple decision thresholds (Table 3a–b), along with RMSE:STDEV ratio of <0.85 and R^2^ of >0.25 indicating a considerable performance by the model (Supplementary Table BC LR1, Supplementary Table BC LR2, Supplementary Table BC DT3 and Supplementary Table BC DT4).

**Table 3a:** Gene predictions at various normalized FPKM thresholds using a Linear Regression model in the Invasive Breast Carcinoma case study

| Invasive Breast Carcinoma: LR model |  |  |
| --- | --- | --- |
| Genes | Normalized FPKM thresholds | AUC |
| CRABP1 | [0.4, 0.8], and 0.3 | 1.0, and 0.97, respectively. |

**Table 3b:** Gene predictions at various normalized FPKM thresholds using a Decision Tree model in the Invasive Breast Carcinoma case study

| Invasive Breast Carcinoma: DT model |  |  |
| --- | --- | --- |
| Genes | Normalized FPKM thresholds | AUC |
| SMTNL2 | [0.4, 0.9], and 0.3 | 1.0, and 0.97, respectively. |

**Table 3c:** Gene predictions at various normalized FPKM thresholds using Multi-Task LASSO and Elastic Net models in the Invasive Breast Carcinoma case study

| Invasive Breast Carcinoma: MTL and MTEN (with prior correlations filtering) |  |  |
| --- | --- | --- |
| Genes | Normalized FPKM thresholds | AUC |
| FABP1 | [0.8, 0.9], and 0.7 | 1.0, and 0.97 respectively. |
| HAS1 | [0.4, 0.9] | 1.0 |

**Table 3d:** Gene predictions at various normalized FPKM thresholds generated using Multi-Task LASSO and Elastic Net models without prior statistical correlation filtering for imaging and gene-FPKM features in the Invasive Breast Carcinoma case study

| Invasive Breast Carcinoma: MTL and MTEN (without prior correlations filtering) |  |  |
| --- | --- | --- |
| Genes | Normalized FPKM thresholds | AUC |
| FAM163A | [0.7, 0.9], and 0.6 | 1.0, and 0.97 respectively. |
| DSG1 | [0.6, 0.9], and 0.5 | 1.0, and 0.94 respectively. |
| SMTNL2 | [0.5, 0.9], and 0.4 | 1.0, and 0.94 respectively. |

The MTL and MTEN models yielded test MSE values of 0.42 and 0.33 respectively, and respective k-fold cross-validation scores of 1.23 and 1.36 (Supplementary reports BC 3-4) during training. FABP1 and HAS1 genes showed high AUC values in the range of [0.97, 1.0] at multiple decision thresholds (Table 3c, Supplementary Table BC MTL6, and Supplementary Table BC MTEN8), along with RMSE: STDEV ratio of <0.85, and R^2^ of >0.25 indicating a considerable performance by the models (Supplementary Table BC MTL5, and Supplementary Table BC MTEN7).

The MTL and MTEN models were trained and tested without carrying out a statistical correlation step (i.e., setting correlation co-efficient threshold as ‘-1.0’ and p-value correction method as ‘none’) and were found to exhibit K-fold cross-validation scores of 1.09 and 1.07, respectively (Supplementary Report BC 5-6) during the training, indicating less overfitting of data by them. MTL identified 3 additional genes (i.e., FAM163A, DSG1 and SMTNL2) having AUC values in the range of [0.94, 1.0] at multiple decision thresholds (Table 3d and Supplementary Table BC MTL10), all of them having RMSE:STDEV ratio of ≤0.85, and R^2^ of >0.25, depicting a considerable model-performance (Supplementary Table BC MTL9), whereas MTEN identified none of the genes fitting that criteria (Supplementary Table BC MTEN11-and Supplementary Table BC MTEN12).

Further assessment of the roles of the identified genes revealed that CRABP1, in conjunction with CRABP2 and FABP1, plays a key role in breast cancer cell responses to the retinoic acid (RA) pathway (R.-Z. Liu et al. 2015). Elevated CRABP1 levels are associated with poor prognosis, high Ki67 immunoreactivity, and high tumor grade in breast cancer (R.-Z. Liu et al. 2015). SMTNL2 is downregulated in breast cancer (Gálvez-Santisteban et al. 2012). HAS1 plays a role in regulating the growth and development of breast tumors and contributes to the development of intratumoral heterogeneity that mimics a cancer stem cell-like phenotype. Its overexpression induces cancer-associated anomalies such as epithelial depletion, chromosomal abnormalities, and micronucleation, thereby demonstrating its role in cancer initiation and progression (Nguyen et al. 2017).

FAM163A regulates the ERK signaling pathway and promotes cell proliferation in squamous cell lung carcinoma; however, its role in IBC has not yet been explored (N. Liu et al. 2019). DSG1 belongs to a family of desmoglein proteins that form a part of the desmosomal complex and that is a known hallmark of cancer (Huber and Petersen 2015). It is a potential biomarker for anal cancer (Myklebust et al. 2012) and a novel target for cancer invasion- and metastasis-focused therapies (Valenzuela-Iglesias et al. 2019). Such literature evidences combined with evidence from ImaGene would encourage researchers to explore the role of these genes in IBC.

In summary, ImaGene facilitates the systematic and user-controlled imaging-based prediction and classification of gene expression in IBC, demonstrating the capability of this platform to identify significant associations between imaging and omics features using correlation analysis as well as AI models in an algorithmic manner. It also allows complete control over multiple operational parameters, ultimately producing a transparent radiogenomics report along with all supporting results files necessary for user interpretation.

### Case Study 2: Head and Neck Cancer

The TCGA Head and Neck Squamous Cell Carcinoma (HNSCC) cohort was accessed to download 540 radiomics features from CT scans of 106 HNSCC patients from a previous imaging study conducted at Stanford University (Mukherjee et al. 2020). Consistent with the results of Case Study 1, the gene expression data (FPKM files) for the entire cohort were downloaded and genes that exhibited FPKM values of greater than or equal to 5 in at least 30 patients were selected for further processing.

The radiomics and gene-FPKM features from 106 samples were used for the experimental run along with the following parameter settings: a) the absolute Pearson-based correlation coefficient threshold was set to 0.7 for linear regression modeling and 0.6 for other model types, b) the p-value correction method was set to Benjamin-Hochberg (BH), and c) the corrected p-value threshold was set to 0.05. Radiomics features were set as data and gene-FPKM values were set as labels. The feature normalization method was set to StandScaler for both inputs, the test size was set to 0.2 (i.e. 20% of the dataset), and the K-fold cross validation splitter (cv) was set to 3.

At the end of the experimental run, a comprehensive radiogenomics report was obtained for each model type (Supplementary reports HNSCC 3-8). These reports presented significant correlations between 12 normalized radiomics features (pertaining to texture, shape, size, and wavelet category) and normalized FPKM values for 21 genes (Supplementary reports HNSCC 3-4). When the correlation coefficient threshold was relaxed to 0.6, 29 additional radiomics features in the wavelet category and 33 additional genes appeared on the list of significant correlations. This expanded feature set was found to improve the training of the DT model.

The LR and DT models yielded test MSE values of 0.57 and 2.1, respectively. For both these models, ImaGene measured AUC values for each gene label across their predicted absolute normalized FPKM decision thresholds in the range of [0,1] (Table 4a–b, Supplementary Table HNSCC DT7, and Supplementary Table HNSCC LR5).

**Table 4a:** Gene predictions at various FPKM thresholds using a Decision Tree model in the HNSCC case study

| HNSCC: DT model |  |  |
| --- | --- | --- |
| Genes | Normalized FPKM thresholds | AUC |
| CEACAM6 | [0.6, 0.9] | 0.97 |

**Table 4b:** Gene predictions at various normalized FPKM thresholds using Multi-Task LASSO and Elastic Net models in the HNSCC case study

| HNSCC: MTEN (with prior correlations) |  |  |
| --- | --- | --- |
| Genes | Normalized FPKM thresholds | AUC |
| NANOG | 0.5 | 1.0 |

**Table 4c:** Gene predictions at various normalized FPKM thresholds using Multi-Task LASSO and Elastic Net models without a prior statistical correlation filtering step for imaging and gene-FPKM feature data in the HNSCC case study

| HNSCC: MTL and MTEN (without prior correlation) |  |  |
| --- | --- | --- |
| Genes | Normalized FPKM thresholds | AUC |
| ACSM2B | [0.8, 0.9], and 0.7 | 1.0, and 0.95 respectively. |
| UPK2 | 0.7 | 0.97 |

LR model demonstrated no reliable association of radiomics features with gene expressions (Supplementary Table HNSCC LR5, and Supplementary Table HNSCC LR6), whereas the DT model (Table 4a) demonstrated the association of radiomics features with high expression of CEACAM6 gene (at AUC=~0.98 at normalized FPKM thresholds of 0.6 to 0.9) with RMSE:Stdev ratio of <0.85 and an R^2^ of >0.25, respectively indicating a considerable model-performance (Supplementary Table HNSCC DT7). A previous study has demonstrated that the focal overexpression of CEACAM6 can promote enhanced head and neck cancer tumorigenesis by suppressing apoptosis (Cameron et al. 2012).

The MTL and MTEN models yielded MSE values of 1.52 and 0.85, respectively, for the test dataset with respective k-fold cross-validation scores of 0.93 and 1.21 (Supplementary reports HNSCC 5-6). MTL model did not yield any reliable associations of radiomics features with gene expressions, whereas MTEN yielded high AUC of 1.0 for NANOG gene at normalized FPKM threshold of 0.5 (Table 4b, Supplementary Table HNSCC MTL11 and Supplementary Table HNSCC MTEN12). NANOG had RMSE:STDEV ratio of <=0.85 and R^2^ of >0.25 making its prediction by MTEN model reliable (Supplementary Table HNSCC MTL11a and Supplementary Table HNSCC MTEN12a). A recent study has shown that NANOG protein expression plays an important role in HNSCC, thereby indicating its potential to serve as a prognostic indicator for patients with HNSCC tumors (Pedregal-Mallo et al. 2020).

The MTL and MTEN models were trained and tested without carrying out the statistical correlation step, leading to the identification of 1 additional gene (i.e., VSIG2) by MTL and 2 additional genes (i.e. ACSM2B and UPK2) by MTEN with AUC values in the range of [0.95, 0.1] at various decision thresholds (Table 4c, Supplementary Table HNSCC MTL9, and Supplementary Table HNSCC MTEN10) along with the RMSE:STDEV ratio of < 0.85 and R^2^ of > 0.25 indicating considerable performance by the models (Supplementary Table HNSCC MTL9a and Supplementary Table HNSCC MTEN10a). Both these models exhibited respective K-fold crossvalidation scores of 1.26 and 1.30 (Supplementary Report HNSCC 7-8) indicating low overfitting of the data by them.

ACSM2B is a known tumor suppressor gene that is regulated by the microRNA miR-31-3p in HNSCC (Oshima et al. 2021). A recent study proposed the usage of UPK2 as one of the top gene signatures to assess the prognostic risk in HNSCC patients (J. Wang et al. 2020).

Overall, these two case studies highlight many important genes that may contribute to IBC and HNSCC incidence and/or progression, thus illustrating the capability of ImaGene as a tool for identifying meaningful associations between imaging features and gene expression data in tumor patients using correlation analysis in conjunction with AI techniques.

## DISCUSSION

ImaGene is a web-based platform that uses a single traceable algorithm to perform statistical correlations and AI modeling techniques as a means of analyzing tumor imaging data. It enables the rapid screening of significantly correlated radiographic and omics features for particular tumors, as well as the construction and testing of different types of AI models using these features as a means of aiding in the prediction of biologically relevant omics features based upon imaging findings. This platform provides superior control over radiogenomic experiments by hosting multiple configuration parameters that can be defined by end users or left at default values. It also obeys FAIR principles, allowing users to safely store and track their experimental data files and to download reports using links available on the job tracking page. The reports are comprehensive HTML documents that provide the outputs of each step of the algorithm, making it easy for users to track the processing of the imaging and omics features through statistical tests and AI modeling. These reports highlight the transparency and easy interpretability of the AI models. The hierarchically clustered heat maps given in these reports represent univariate, bivariate, and multivariate statistical correlations between the imaging and omics features, and the MSE, RMSE: STDEV, and AUC values reflect the reliability of the AI models when predicting and classifying omics labels from imaging data. Thus, these reports prepared by the platform provide a comprehensive overview of the entire analytical process and permit users to interpret their datasets while providing with the experimental parameters necessary to arrive at meaningful conclusions.

ImaGene aims to advance current AI modeling through its flexibility, ease of interpretation, and user-friendly design, ensuring that users can confidently design and interpret their radiogenomic experiments. Relative to other radiogenomic applications such as Imaging-Amaretto, Imaging-Community Amaretto, and Musa, ImaGene’s features have the potential to make it widely adopted in hospitals and laboratories worldwide as a tool for testing associations between imaging and omics data for different tumor types, paving the way for significant discoveries in the fields of radiogenomics and oncological research.

As a proof of concept, we tested the performance of ImaGene for the prediction and classification of omics profiles from imaging features using IBC and HNSCC datasets. We identified strong associations between the expression of 6 genes (i.e., CRABP1, SMTNL2, FABP1, HAS1, FAM163A and DSG1) and imaging features such as tumor size, shape, enhancement, and kinetic curve assessments in IBC (Supplementary Report BC 1-6 and Tables:3a–3d). Further research into the roles of these genes in cancer revealed that *all* of them have been shown to play an indirect or direct role in shaping IBC tumorigenesis. In our HNSCC case study, ImaGene further detected strong associations between the expression of 4 genes (i.e., CEACAM6, NANOG, ACSM2B, and UPK2) and imaging attributes such as tumor texture, shape, size, and wavelet features (Supplementary Report HNSCC 3-8 and Tables:4a-4d). All of these genes have also been previously demonstrated to be associated with therapeutic outcomes and survival among HNSCC patients.

ImaGene can facilitate collaborations between researchers, making it the sharing of radiogenomics reports more convenient and ultimately leading to the advancement of research in the cancer and precision medicine space. Our case studies have sought to provide a basis for the radiogenomic analysis through the processing of imaging and omics feature data in two tumor types while permitting robust control over experimental parameters. We have successfully demonstrated the configuration and usage of statistical correlations and AI modeling on our platform, leading to the construction of reliable AI models that can predict biologically relevant omics features from imaging data. Analysis of multiple datasets of different types and subtypes of tumors in ImaGene can contribute to the development of a radiogenomic knowledge-base that can be used in cancer research, drug discovery, and precision medicine, allowing the outcomes of this platform to undergo rapid translation for clinical use in the near future.

## Supporting information

Supplementary_table_HNSCC_MTEN_10

Supplementary_table_HNSCC_MTEN_10a

Supplementary_table_HNSCC_MTL_9

Supplementary_table_HNSCC_MTL_9a

Supplementary_table_HNSCC_MTL_11a

Supplementary_table_HNSCC_MTL_11

Supplementary_Table_HNSCC_DT_8

Supplementary_Table_HNSCC_DT_7

Supplementary_Table_HNSCC_LR_5

Supplementary_Table_HNSCC_LR_6

Supplementary_table_BC_MTEN_11

Supplementary_table_BC_MTEN_12

Supplementary_table_BC_MTL_10

Supplementary_table_BC_MTL_9

Supplementary_Table_BC_MTEN_7

Supplementary Table_BC_MTEN_8

Supplementary Table_BC_MTL_6

Supplementary_Table_BC_MTL_5

Supplementary_Table_HNSCC_MTEN_12a

Supplementary_Table_HNSCC_MTEN_12

Supplementary_report_BC_3

Supplementary_table_BC-Gene_FPKM

Supplementary_table_BC-Radiomic_features

Supplementary_report_BC_4

Supplementary_Report_BC_6

Supplementary_Report_BC_5

Supplementary_Report_HNSCC_6

Supplementary_Report_HNSCC_5

Supplementary_table_HNSCC-Radiomic_features

Supplementary_table_HNSCC-Gene_FPKM

Supplementary_Report_HNSCC_8

Supplementary_Report_HNSCC_7

Supplementary_report_HNSCC_4

Supplementary_report_HNSCC_3

Supplementary_report_BC_2

Supplementary_report_BC_1

