## Supplementary_Report_BC_6 for "ImaGene: A web-based software platform for tumor radiogenomic evaluation and reporting"

### Radiogenomics Analysis Report

##### 23/09/2021 14:59:41

#### ----------------------------Model inputs-------------------------------

##### Mode:Train

##### Model:multiTaskLinearModel

##### Params:default

##### No. of imaging features provided: 36

##### No. of gene features provided:976

##### SampleID check results: 'The SampleIDs match for imaging and gene features'

##### No. of samples: 89

##### performing Stand\_scaler normalization for imaging features

##### performing Stand\_scaler for gene features

#### --------------------------Multivariate Correlations (pearson based)-----------------------


### ----------------------------------Features with highly significant correlations-------------------------------------

#### Below is the list of imaging features

###### ['Contrast\_(T1)', 'Correlation\_(T2)', 'Curve\_shape\_index\_(K5)', 'Difference\_Entropy\_(T3)', 'Difference\_Variance\_(T4)', 'E1\_(K6)', 'Effective\_Diameter\_(S2)', 'Energy\_(T5)', 'Enhancement-Variance\_Time\_to\_Peak\_(E2)', 'Enhancement-variance\_Decreasing\_Rate\_(E4)', 'Enhancement-variance\_Increasing\_Rate\_(E3)', 'Entropy\_(T6)', 'Homogeneity\_(T7)', 'IMC1\_(T8)', 'IMC2\_(T9)', 'Irregularity\_(G2)', 'Margin\_Sharpness\_(M1)', 'Maximum\_Correlation\_Coefficient\_(T10)', 'Maximum\_Diameter\_(S5)', 'Maximum\_enhancement-variance\_(E1)', 'Maximum\_enhancement\_(K1)', 'Signal\_Enhancement\_Ratio\_(SER)\_(K7)', 'Size-Lesion\_volume\_(S1)', 'Sphericity\_(G1)', 'Sum\_Average\_(T11)', 'Sum\_Entropy\_(T12)', 'Sum\_Variance\_(T13)', 'Surface\_Area\_(S3)', 'Surface\_Area\_to\_Volume\_ratio\_(G3)', 'Time\_to\_peak\_(K2)', 'Uptake\_rate\_(K3)', 'Variance\_(T14)', 'Variance\_of\_Margin\_Sharpness\_(M2)', 'Variance\_of\_Radial\_Gradient\_Histogram\_(vRGH)\_(M3)', 'Volume\_of\_most\_enhancing\_voxels\_(S4)', 'Washout\_rate\_(K4)']

#### Below is the list of gene features

###### ['ABCB5', 'AC234582.1', 'ACSM2B', 'ACTG2', 'ADH1A', 'ADRA2C', 'ADRB1', 'AGT', 'AGXT', 'AHSP', 'AICDA', 'AIM2', 'AL138751.1', 'ALPPL2', 'AMER2', 'AMHR2', 'AMPD1', 'ANXA10', 'ANXA13', 'ANXA8', 'AOC1', 'APOA2', 'APOB', 'APOC3', 'APOD', 'APOH', 'AR', 'ARG1', 'ARPP21', 'ASCL2', 'ASIC4', 'ATCAY', 'ATP1A3', 'ATP6V0D2', 'ATP6V1G3', 'AXIN2', 'AZGP1', 'BBOX1', 'BCHE', 'BEX1', 'BHLHA15', 'BHMT', 'BHMT2', 'BLK', 'BMP6', 'BMPER', 'BNC1', 'BPI', 'BRINP2', 'BSND', 'BTK', 'C10orf99', 'C11orf21', 'C16orf89', 'C18orf42', 'C1QTNF4', 'C1orf228', 'C1orf61', 'C2orf88', 'C4BPA', 'C7', 'C8B', 'C8orf46', 'CA14', 'CA9', 'CACNG7', 'CALB2', 'CALML5', 'CAPS', 'CCL13', 'CCL20', 'CCL25', 'CCNA1', 'CCR2', 'CCR9', 'CD177', 'CD19', 'CD1B', 'CD1C', 'CD1E', 'CD22', 'CD3D', 'CD70', 'CD79B', 'CD8B', 'CDH15', 'CDHR5', 'CDK5R2', 'CDO1', 'CDX1', 'CEACAM3', 'CES1', 'CFHR1', 'CFTR', 'CHAD', 'CHGA', 'CHRNA3', 'CHST4', 'CILP', 'CLC', 'CLCNKB', 'CLDN10', 'CLDN16', 'CLDN6', 'CLEC17A', 'CLECL1', 'CLRN3', 'CNTFR', 'COMP', 'CORO2B', 'CPA4', 'CPB1', 'CPB2', 'CPLX2', 'CPNE5', 'CRABP1', 'CRISP3', 'CRP', 'CRYAB', 'CST1', 'CTNND2', 'CTSE', 'CTSV', 'CXCL8', 'CXCR2', 'CYP11A1', 'CYP11B1', 'CYP17A1', 'CYP4B1', 'CYSLTR1', 'DAAM2', 'DACH1', 'DAPL1', 'DBH', 'DCDC2', 'DCX', 'DDC', 'DEFA3', 'DEFA4', 'DEFB1', 'DERL3', 'DES', 'DKK1', 'DLX3', 'DLX5', 'DMGDH', 'DMRT2', 'DNER', 'DPEP1', 'DPPA4', 'DPPA5', 'DRD2', 'DSC3', 'DSG1', 'DSG3', 'DUSP5', 'EDNRB', 'EEF1A2', 'EFHD1', 'ELANE', 'ELAVL3', 'EMR3', 'ENPEP', 'EPHX3', 'EPS8L3', 'ESR1', 'EYA1', 'F13B', 'F2', 'F7', 'FABP1', 'FABP7', 'FAIM3', 'FAM101A', 'FAM129C', 'FAM163A', 'FAM166B', 'FAM19A3', 'FAM19A4', 'FAM3D', 'FAM43B', 'FAT2', 'FCN1', 'FCRL2', 'FCRL3', 'FCRL5', 'FCRLA', 'FGF18', 'FGF8', 'FGF9', 'FLT3', 'FOLH1', 'FOXA1', 'FOXC1', 'FOXD1', 'FOXE1', 'FOXJ1', 'FOXN1', 'G0S2', 'G6PC', 'GALNT9', 'GAPDHS', 'GAPT', 'GATA3', 'GATA4', 'GATA6', 'GBP6', 'GCG', 'GCGR', 'GDNF', 'GFAP', 'GFRA1', 'GFRA3', 'GIP', 'GJB1', 'GLYAT', 'GNG3', 'GNG7', 'GPA33', 'GPM6B', 'GPR64', 'GPRC5B', 'GPRC5D', 'GYG2', 'GZMK', 'HABP2', 'HAND1', 'HAND2', 'HAS1', 'HEPACAM', 'HEPACAM2', 'HIST1H1B', 'HIST1H1D', 'HIST1H2AB', 'HIST1H2AH', 'HIST1H2AI', 'HIST1H2AJ', 'HIST1H2AM', 'HIST1H2BG', 'HIST1H2BH', 'HIST1H2BL', 'HIST1H2BO', 'HIST1H3B', 'HIST1H3D', 'HIST1H3F', 'HIST1H3G', 'HIST1H3J', 'HIST1H4C', 'HIST1H4D', 'HIST1H4L', 'HIST2H2AC', 'HLA-DOB', 'HMP19', 'HNF1B', 'HNF4A', 'HOGA1', 'HOXA5', 'HOXB13', 'HOXB6', 'HOXC10', 'HOXD11', 'HOXD8', 'HP', 'HPX', 'HR', 'HRG', 'HSD3B2', 'HSPB2', 'HSPB6', 'ICAM3', 'IFITM5', 'IGF2', 'IGFBP1', 'IGJ', 'IGLL1', 'IL18RAP', 'IL21R', 'IL36G', 'IL36RN', 'INS', 'INSC', 'INSM2', 'IRF4', 'ISL1', 'ITIH1', 'ITIH5', 'ITLN2', 'KCNJ16', 'KCNK15', 'KCNQ2', 'KHDC3L', 'KIF26A', 'KL', 'KLHDC8A', 'KLK1', 'KLK4', 'KLK5', 'KLK6', 'KLK7', 'KLK8', 'KNG1', 'KREMEN2', 'KRT13', 'KRT16', 'KRT17', 'KRT20', 'KRT5', 'KRT6A', 'KRT6B', 'KRTDAP', 'LAMP5', 'LCN12', 'LEFTY1', 'LGALS4', 'LGR5', 'LIN28A', 'LMX1B', 'LRMP', 'LRRC14B', 'LRRC26', 'LTB', 'LYG1', 'LYPD6B', 'LZTS1', 'MAB21L1', 'MAB21L2', 'MAG', 'MAL', 'MAT1A', 'MB', 'MECOM', 'MEF2B', 'MEOX1', 'MEP1A', 'MFAP5', 'MGAM', 'MGP', 'MIA', 'MIXL1', 'MLANA', 'MLC1', 'MLPH', 'MMP1', 'MMP12', 'MMP17', 'MMP25', 'MMP9', 'MRGPRF', 'MS4A1', 'MS4A3', 'MSMB', 'MT1H', 'MTRNR2L12', 'MTRNR2L8', 'MUC13', 'MUC15', 'MUC3A', 'MXRA8', 'NANOG', 'NANOGP8', 'NAPSA', 'NAT8', 'NCAN', 'NDNF', 'NDP', 'NDUFA4L2', 'NEFL', 'NEFM', 'NKX2-1', 'NKX3-1', 'NLRP7', 'NNAT', 'NOX1', 'NPW', 'NPY', 'NR1H4', 'NTRK1', 'NTS', 'OCA2', 'OR51E1', 'ORM1', 'ORM2', 'P2RX1', 'P2RX3', 'P2RX5', 'PADI4', 'PAX2', 'PAX5', 'PAX9', 'PCDH18', 'PEBP4', 'PEG10', 'PENK', 'PGLYRP1', 'PHGR1', 'PHOX2B', 'PIGR', 'PKP1', 'PLA2G2A', 'PLA2G2D', 'PLAC8', 'PLG', 'PLP1', 'PMEL', 'PNCK', 'PNMAL2', 'PNOC', 'POSTN', 'POU2AF1', 'POU2F2', 'PPP1R1A', 'PRAC2', 'PRDM14', 'PRG4', 'PRPH', 'PRSS57', 'PTCRA', 'PTGDS', 'PTGER3', 'PTHLH', 'PTPRN', 'PTPRN2', 'PTPRZ1', 'PVALB', 'RAB37', 'RAB38', 'RAB3B', 'RAB3C', 'RAB44', 'RASGRP3', 'RBP4', 'RBP5', 'REEP6', 'REG1A', 'REG4', 'RGL4', 'RGS20', 'RHCG', 'RNASE2', 'RNASE3', 'RNF186', 'RSPO1', 'RTN1', 'RUNDC3A', 'RXRG', 'S100A1', 'S100A14', 'S100B', 'S100P', 'SAA1', 'SCARF2', 'SCG5', 'SCGB1A1', 'SCRG1', 'SDR16C5', 'SERPINA5', 'SERPINA6', 'SERPINB10', 'SERPINC1', 'SERPINE2', 'SFRP2', 'SFTA3', 'SFTPA1', 'SFTPA2', 'SFTPB', 'SFTPD', 'SH2D1A', 'SHISA2', 'SHISA3', 'SHISA8', 'SLAMF7', 'SLC18A1', 'SLC22A2', 'SLC22A31', 'SLC24A2', 'SLC26A4', 'SLC26A7', 'SLC27A6', 'SLC28A1', 'SLC30A2', 'SLC3A1', 'SLC45A3', 'SLC4A1', 'SLC4A4', 'SLC6A17', 'SLC7A4', 'SLC7A8', 'SMTNL2', 'SNAP91', 'SNCA', 'SNCG', 'SNX22', 'SOX1', 'SOX17', 'SOX18', 'SOX8', 'SP7', 'SPDEF', 'SPIB', 'SPINK1', 'SPINK2', 'SPNS3', 'SPOCK3', 'SPON1', 'SPRR3', 'SPX', 'ST6GALNAC1', 'STAR', 'STC2', 'STEAP2', 'STK32A', 'STMN2', 'STMN4', 'SYP', 'TBX3', 'TCERG1L', 'TCF21', 'TCL1A', 'TCN1', 'TFR2', 'TGM1', 'TH', 'TLR10', 'TLX2', 'TM4SF4', 'TMEM100', 'TMEM119', 'TMEM151A', 'TMEM179', 'TMEM213', 'TMEM238', 'TMEM252', 'TMPRSS11D', 'TNF', 'TNFRSF13B', 'TNFRSF13C', 'TNFRSF17', 'TNR', 'TP63', 'TRABD2A', 'TRABD2B', 'TRIM31', 'TRPS1', 'TRPV6', 'TSPAN32', 'TSPAN8', 'TTR', 'TUBB2B', 'TUBB4A', 'TXNDC5', 'UGT2B4', 'UPK2', 'UPK3A', 'UPK3B', 'UTF1', 'UTS2R', 'VENTX', 'VGLL1', 'VPREB1', 'VRTN', 'VSTM1', 'VSTM2A', 'VTN', 'ZBED2', 'ZCCHC12']

#### Number of imaging features

### 36

#### Number of gene features

### 565

### -------------------------------Number of Samples for Training and Testing---------------------------------

##### No. of samples for training:71

##### No. of samples for test:18

#### --------------------------Model Summary-----------------------

##### Model Type : multiTaskLinearModel

###### Cross Validation Metrics:

###### Parameters: cv=3 scoring=neg\_mean\_squared\_error

###### Cross validation score:1.07449439385

##### Model Parameters:

###### normalize:False

###### warm\_start:False

###### selection:cyclic

###### fit\_intercept:True

###### l1\_ratio:0.5

###### max\_iter:1000

###### random\_state:None

###### tol:0.0001

###### copy\_X:True

###### alpha:1.0

#### ----------------Model evaluation for Train data--------------------

##### Min Square Error for the Model

###### MSE of train\_eval set:0.717258202058

###### No. of features showing LOW 'RMSE/Stdev' (<=1.0): 565

###### All such features with their Low 'RMSE/Stdev' values could be found in output file: train\_eval\_multiTaskLinearModel\_Labels\_with\_Low\_Ratio.csv No. of features showing HIGH 'RMSE/Stdev' (>1.0): 0 All such features with their High 'RMSE/Stdev' values could be found in output file: train\_eval\_multiTaskLinearModel\_Labels\_with\_High\_Ratio.csv Model evaluation for Train data for label features showing Low 'RMSE/Stdev' (<=1.0) Content-type: text/html Content-type: text/html ----------------Model evaluation for Test data-------------------- Min Square Error for the Model MSE of test\_eval set:1.24768165209 No. of features showing LOW 'RMSE/Stdev' (<=1.0): 103 All such features with their Low 'RMSE/Stdev' values could be found in output file: test\_eval\_multiTaskLinearModel\_Labels\_with\_Low\_Ratio.csv No. of features showing HIGH 'RMSE/Stdev' (>1.0): 460 All such features with their High 'RMSE/Stdev' values could be found in output file: test\_eval\_multiTaskLinearModel\_Labels\_with\_High\_Ratio.csv Model evaluation for Test data for label features showing Low 'RMSE/Stdev' (<=1.0) Content-type: text/html Content-type: text/html
