## Supplementary_Report_BC_5 for "ImaGene: A web-based software platform for tumor radiogenomic evaluation and reporting"

###### Parameters: cv=3 scoring=neg\_mean\_squared\_error

###### Cross validation score:1.09212395238

#### Model Parameters:

##### normalize:False

##### warm\_start:False

##### selection:cyclic

##### fit\_intercept:True

##### max\_iter:1000

##### random\_state:None

##### tol:0.0001

##### copy\_X:True

##### alpha:1.0

### ----------------Model evaluation for Train data--------------------

#### Min Square Error for the Model

##### MSE of train\_eval set:0.732274624443

##### No. of features showing LOW 'RMSE/Stdev' (<=1.0): 565

###### All such features with their Low 'RMSE/Stdev' values could be found in output file: train\_eval\_multiTaskLASSO\_Labels\_with\_Low\_Ratio.csv No. of features showing HIGH 'RMSE/Stdev' (>1.0): 0 All such features with their High 'RMSE/Stdev' values could be found in output file: train\_eval\_multiTaskLASSO\_Labels\_with\_High\_Ratio.csv Model evaluation for Train data for label features showing Low 'RMSE/Stdev' (<=1.0) Content-type: text/html Content-type: text/html ----------------Model evaluation for Test data-------------------- Min Square Error for the Model MSE of test\_eval set:1.27548202487 No. of features showing LOW 'RMSE/Stdev' (<=1.0): 109 All such features with their Low 'RMSE/Stdev' values could be found in output file: test\_eval\_multiTaskLASSO\_Labels\_with\_Low\_Ratio.csv No. of features showing HIGH 'RMSE/Stdev' (>1.0): 456 All such features with their High 'RMSE/Stdev' values could be found in output file: test\_eval\_multiTaskLASSO\_Labels\_with\_High\_Ratio.csv Model evaluation for Test data for label features showing Low 'RMSE/Stdev' (<=1.0) Content-type: text/html Content-type: text/html
