## Supplementary_Report_HNSCC_6 for "ImaGene: A web-based software platform for tumor radiogenomic evaluation and reporting"

### Radiogenomics Analysis Report

##### 22/09/2021 17:01:46

#### ----------------------------Model inputs-------------------------------

##### Mode:Train

##### Model:multiTaskLinearModel

##### Params:default

##### No. of imaging features provided: 540

##### No. of gene features provided:976

##### SampleID check results: 'The SampleIDs match for imaging and gene features'

##### No. of samples: 106

##### performing Stand\_scaler normalization for imaging features

##### performing Stand\_scaler for gene features

#### --------------------------Multivariate Correlations (pearson based)-----------------------


### ----------------------------------Features with highly significant correlations-------------------------------------

#### Below is the list of imaging features

###### ['GLRLM\_LongRunEmphasis', 'GLRLM\_LongRunLowGrayLevelEmphasis', 'GLRLM\_RunLengthNonUniformity', 'GLSZM\_LZE', 'GLSZM\_LZHGE', 'GLSZM\_LZLGE', 'globalHistogram\_Mode', 'shapeSize\_Compactness2', 'shapeSize\_SurfaceArea', 'waveletHHH\_GLRLM\_LongRunHighGrayLevelEmphasis', 'waveletHHH\_GLRLM\_LongRunLowGrayLevelEmphasis', 'waveletHHH\_GLRLM\_RunLengthNonUniformity', 'waveletHHL\_GLRLM\_LongRunLowGrayLevelEmphasis', 'waveletHHL\_firstOrder\_QuartileCoefficientDispersion', 'waveletHLH\_GLCM\_Energy', 'waveletHLH\_GLRLM\_LongRunEmphasis', 'waveletHLH\_GLRLM\_LongRunHighGrayLevelEmphasis', 'waveletHLH\_GLRLM\_RunLengthNonUniformity', 'waveletHLH\_firstOrder\_CoefficientVariation', 'waveletHLL\_GLRLM\_GrayLevelNonUniformity', 'waveletHLL\_GLRLM\_LongRunHighGrayLevelEmphasis', 'waveletHLL\_GLRLM\_RunLengthNonUniformity', 'waveletHLL\_firstOrder\_QuartileCoefficientDispersion', 'waveletLHH\_GLRLM\_LongRunHighGrayLevelEmphasis', 'waveletLHH\_GLRLM\_LongRunLowGrayLevelEmphasis', 'waveletLHH\_GLRLM\_RunLengthNonUniformity', 'waveletLHL\_GLRLM\_GrayLevelNonUniformity', 'waveletLHL\_firstOrder\_QuartileCoefficientDispersion', 'waveletLLH\_GLRLM\_GrayLevelNonUniformity', 'waveletLLH\_GLRLM\_LongRunEmphasis', 'waveletLLH\_GLRLM\_LongRunHighGrayLevelEmphasis', 'waveletLLH\_GLRLM\_RunLengthNonUniformity', 'waveletLLH\_firstOrder\_CoefficientVariation', 'waveletLLH\_firstOrder\_Mean', 'waveletLLH\_firstOrder\_Median', 'waveletLLL\_GLCM\_Energy', 'waveletLLL\_GLRLM\_GrayLevelNonUniformity', 'waveletLLL\_GLRLM\_LongRunEmphasis', 'waveletLLL\_GLRLM\_LongRunLowGrayLevelEmphasis', 'waveletLLL\_GLRLM\_RunLengthNonUniformity', 'waveletLLL\_firstOrder\_Maximum']

#### Below is the list of gene features

###### ['11-Mar', 'ADRA2C', 'AGXT', 'ATP6V0D2', 'AZGP1', 'BIRC7', 'C18orf42', 'C1QL1', 'C8B', 'CADM3', 'CDH16', 'CEACAM6', 'CUX2', 'CYP17A1', 'DCSTAMP', 'DCT', 'DPPA3', 'EEF1A2', 'FABP7', 'G6PC', 'GATA3', 'GFRA3', 'HIST1H3G', 'HIST1H4D', 'IGLL1', 'INSC', 'ITLN1', 'KLK4', 'LAMP5', 'LIN28A', 'MAL', 'MMP8', 'MMP9', 'MYH11', 'NANOG', 'NPY', 'PANX3', 'PEG3', 'PGLYRP2', 'POU5F1', 'PRDM14', 'PSMB11', 'PVALB', 'SERPINA1', 'SERPINB10', 'SLC22A31', 'SLC26A7', 'SLC3A1', 'SP7', 'STMN2', 'TMEM52B', 'TSHR', 'TYR', 'VTCN1']

#### Number of imaging features

### 41

#### Number of gene features

### 54

### -------------------------------Number of Samples for Training and Testing---------------------------------

##### No. of samples for training:84

##### No. of samples for test:22

#### --------------------------Model Summary-----------------------

##### Model Type : multiTaskLinearModel

###### Cross Validation Metrics:

###### Parameters: cv=3 scoring=neg\_mean\_squared\_error

###### Cross validation score:1.21075627758

##### Model Parameters:

###### normalize:False

###### warm\_start:False

###### selection:cyclic

###### fit\_intercept:True

###### l1\_ratio:0.5

###### max\_iter:1000

###### random\_state:None

###### tol:0.0001

###### copy\_X:True

###### alpha:1.0

#### ----------------Model evaluation for Train data--------------------

##### Min Square Error for the Model

###### MSE of train\_eval set:0.571045581713

###### No. of features showing LOW 'RMSE/Stdev' (<=1.0): 54

###### All such features with their Low 'RMSE/Stdev' values could be found in output file: train\_eval\_multiTaskLinearModel\_Labels\_with\_Low\_Ratio.csv No. of features showing HIGH 'RMSE/Stdev' (>1.0): 0 All such features with their High 'RMSE/Stdev' values could be found in output file: train\_eval\_multiTaskLinearModel\_Labels\_with\_High\_Ratio.csv Model evaluation for Train data for label features showing Low 'RMSE/Stdev' (<=1.0) Content-type: text/html Content-type: text/html ----------------Model evaluation for Test data-------------------- Min Square Error for the Model MSE of test\_eval set:0.850476820968 No. of features showing LOW 'RMSE/Stdev' (<=1.0): 8 All such features with their Low 'RMSE/Stdev' values could be found in output file: test\_eval\_multiTaskLinearModel\_Labels\_with\_Low\_Ratio.csv No. of features showing HIGH 'RMSE/Stdev' (>1.0): 46 All such features with their High 'RMSE/Stdev' values could be found in output file: test\_eval\_multiTaskLinearModel\_Labels\_with\_High\_Ratio.csv Model evaluation for Test data for label features showing Low 'RMSE/Stdev' (<=1.0) Content-type: text/html Content-type: text/html
