## Supplementary_Report_HNSCC_8 for "ImaGene: A web-based software platform for tumor radiogenomic evaluation and reporting"

### Radiogenomics Analysis Report

##### 22/09/2021 13:01:03

#### ----------------------------Model inputs-------------------------------

##### Mode:Train

##### Model:multiTaskLinearModel

##### Params:default

##### No. of imaging features provided: 540

##### No. of gene features provided:976

##### SampleID check results: 'The SampleIDs match for imaging and gene features'

##### No. of samples: 106

##### performing Stand\_scaler normalization for imaging features

##### performing Stand\_scaler for gene features

#### --------------------------Multivariate Correlations (pearson based)-----------------------


### ----------------------------------Features with highly significant correlations-------------------------------------

#### Below is the list of imaging features

###### ['GLCM\_Autocorrelation', 'GLCM\_ClusterProminence', 'GLCM\_ClusterShade', 'GLCM\_ClusterTendency', 'GLCM\_Contrast', 'GLCM\_Correlation', 'GLCM\_DifferenceEntropy', 'GLCM\_DifferenceVariance', 'GLCM\_Dissimilarity', 'GLCM\_Energy', 'GLCM\_Entropy', 'GLCM\_Homogeneity1', 'GLCM\_Homogeneity2', 'GLCM\_IDMN', 'GLCM\_IDN', 'GLCM\_IMC1', 'GLCM\_IMC2', 'GLCM\_InverseVariance', 'GLCM\_MaximumProbability', 'GLCM\_SumAverage', 'GLCM\_SumEntropy', 'GLCM\_SumVariance', 'GLCM\_Variance', 'GLRLM\_GrayLevelNonUniformity', 'GLRLM\_HighGrayLevelRunEmphasis', 'GLRLM\_LongRunEmphasis', 'GLRLM\_LongRunHighGrayLevelEmphasis', 'GLRLM\_LongRunLowGrayLevelEmphasis', 'GLRLM\_LowGrayLevelRunEmphasis', 'GLRLM\_RunLengthNonUniformity', 'GLRLM\_RunPercentage', 'GLRLM\_ShortRunEmphasis', 'GLRLM\_ShortRunHighGrayLevelEmphasis', 'GLRLM\_ShortRunLowGrayLevelEmphasis', 'GLSZM\_GLN', 'GLSZM\_GLV', 'GLSZM\_HGZE', 'GLSZM\_LGZE', 'GLSZM\_LZE', 'GLSZM\_LZHGE', 'GLSZM\_LZLGE', 'GLSZM\_SZE', 'GLSZM\_SZHGE', 'GLSZM\_SZLGE', 'GLSZM\_ZP', 'GLSZM\_ZSN', 'GLSZM\_ZSV', 'NGTDM\_Busyness', 'NGTDM\_Coarseness', 'NGTDM\_Complexity', 'NGTDM\_Contrast', 'NGTDM\_Strength', 'firstOrder\_CoefficientVariation', 'firstOrder\_Energy', 'firstOrder\_Entropy', 'firstOrder\_InterquartileRange', 'firstOrder\_Kurtosis', 'firstOrder\_Maximum', 'firstOrder\_Mean', 'firstOrder\_MeanAbsoluteDeviation', 'firstOrder\_Median', 'firstOrder\_MedianAbsoluteDeviation', 'firstOrder\_Minimum', 'firstOrder\_Percentile10', 'firstOrder\_Percentile90', 'firstOrder\_QuartileCoefficientDispersion', 'firstOrder\_Range', 'firstOrder\_RobustMeanAbsoluteDeviation', 'firstOrder\_RootMeanSquare', 'firstOrder\_Skewness', 'firstOrder\_StandardDeviation', 'firstOrder\_Uniformity', 'firstOrder\_Variance', 'globalHistogram\_CoefficientVariation', 'globalHistogram\_Entropy', 'globalHistogram\_InterquartileRange', 'globalHistogram\_Kurtosis', 'globalHistogram\_MeanAbsoluteDeviation', 'globalHistogram\_Median', 'globalHistogram\_MedianAbsoluteDeviation', 'globalHistogram\_Mode', 'globalHistogram\_Percentile10', 'globalHistogram\_Percentile90', 'globalHistogram\_QuartileCoefficientDispersion', 'globalHistogram\_Range', 'globalHistogram\_RobustMeanAbsoluteDeviation', 'globalHistogram\_Skewness', 'globalHistogram\_Uniformity', 'globalHistogram\_Variance', 'shapeSize\_Compactness1', 'shapeSize\_Compactness2', 'shapeSize\_Eccentricity', 'shapeSize\_Maximum3DDiameter', 'shapeSize\_Solidity', 'shapeSize\_SphericalDisproportion', 'shapeSize\_Sphericity', 'shapeSize\_SurfaceArea', 'shapeSize\_SurfaceVolumeRatio', 'shapeSize\_Volume', 'waveletHHH\_GLCM\_Autocorrelation', 'waveletHHH\_GLCM\_ClusterProminence', 'waveletHHH\_GLCM\_ClusterShade', 'waveletHHH\_GLCM\_ClusterTendency', 'waveletHHH\_GLCM\_Contrast', 'waveletHHH\_GLCM\_Correlation', 'waveletHHH\_GLCM\_DifferenceEntropy', 'waveletHHH\_GLCM\_DifferenceVariance', 'waveletHHH\_GLCM\_Dissimilarity', 'waveletHHH\_GLCM\_Energy', 'waveletHHH\_GLCM\_Entropy', 'waveletHHH\_GLCM\_Homogeneity1', 'waveletHHH\_GLCM\_Homogeneity2', 'waveletHHH\_GLCM\_IDMN', 'waveletHHH\_GLCM\_IDN', 'waveletHHH\_GLCM\_IMC1', 'waveletHHH\_GLCM\_IMC2', 'waveletHHH\_GLCM\_InverseVariance', 'waveletHHH\_GLCM\_MaximumProbability', 'waveletHHH\_GLCM\_SumAverage', 'waveletHHH\_GLCM\_SumEntropy', 'waveletHHH\_GLCM\_SumVariance', 'waveletHHH\_GLCM\_Variance', 'waveletHHH\_GLRLM\_GrayLevelNonUniformity', 'waveletHHH\_GLRLM\_HighGrayLevelRunEmphasis', 'waveletHHH\_GLRLM\_LongRunEmphasis', 'waveletHHH\_GLRLM\_LongRunHighGrayLevelEmphasis', 'waveletHHH\_GLRLM\_LongRunLowGrayLevelEmphasis', 'waveletHHH\_GLRLM\_LowGrayLevelRunEmphasis', 'waveletHHH\_GLRLM\_RunLengthNonUniformity', 'waveletHHH\_GLRLM\_RunPercentage', 'waveletHHH\_GLRLM\_ShortRunEmphasis', 'waveletHHH\_GLRLM\_ShortRunHighGrayLevelEmphasis', 'waveletHHH\_GLRLM\_ShortRunLowGrayLevelEmphasis', 'waveletHHH\_firstOrder\_CoefficientVariation', 'waveletHHH\_firstOrder\_Energy', 'waveletHHH\_firstOrder\_Entropy', 'waveletHHH\_firstOrder\_InterquartileRange', 'waveletHHH\_firstOrder\_Kurtosis', 'waveletHHH\_firstOrder\_Maximum', 'waveletHHH\_firstOrder\_Mean', 'waveletHHH\_firstOrder\_MeanAbsoluteDeviation', 'waveletHHH\_firstOrder\_Median', 'waveletHHH\_firstOrder\_MedianAbsoluteDeviation', 'waveletHHH\_firstOrder\_Minimum', 'waveletHHH\_firstOrder\_Percentile10', 'waveletHHH\_firstOrder\_Percentile90', 'waveletHHH\_firstOrder\_QuartileCoefficientDispersion', 'waveletHHH\_firstOrder\_Range', 'waveletHHH\_firstOrder\_RobustMeanAbsoluteDeviation', 'waveletHHH\_firstOrder\_RootMeanSquare', 'waveletHHH\_firstOrder\_Skewness', 'waveletHHH\_firstOrder\_StandardDeviation', 'waveletHHH\_firstOrder\_Uniformity', 'waveletHHH\_firstOrder\_Variance', 'waveletHHL\_GLCM\_Autocorrelation', 'waveletHHL\_GLCM\_ClusterProminence', 'waveletHHL\_GLCM\_ClusterShade', 'waveletHHL\_GLCM\_ClusterTendency', 'waveletHHL\_GLCM\_Contrast', 'waveletHHL\_GLCM\_Correlation', 'waveletHHL\_GLCM\_DifferenceEntropy', 'waveletHHL\_GLCM\_DifferenceVariance', 'waveletHHL\_GLCM\_Dissimilarity', 'waveletHHL\_GLCM\_Energy', 'waveletHHL\_GLCM\_Entropy', 'waveletHHL\_GLCM\_Homogeneity1', 'waveletHHL\_GLCM\_Homogeneity2', 'waveletHHL\_GLCM\_IDMN', 'waveletHHL\_GLCM\_IDN', 'waveletHHL\_GLCM\_IMC1', 'waveletHHL\_GLCM\_IMC2', 'waveletHHL\_GLCM\_InverseVariance', 'waveletHHL\_GLCM\_MaximumProbability', 'waveletHHL\_GLCM\_SumAverage', 'waveletHHL\_GLCM\_SumEntropy', 'waveletHHL\_GLCM\_SumVariance', 'waveletHHL\_GLCM\_Variance', 'waveletHHL\_GLRLM\_GrayLevelNonUniformity', 'waveletHHL\_GLRLM\_HighGrayLevelRunEmphasis', 'waveletHHL\_GLRLM\_LongRunEmphasis', 'waveletHHL\_GLRLM\_LongRunHighGrayLevelEmphasis', 'waveletHHL\_GLRLM\_LongRunLowGrayLevelEmphasis', 'waveletHHL\_GLRLM\_LowGrayLevelRunEmphasis', 'waveletHHL\_GLRLM\_RunLengthNonUniformity', 'waveletHHL\_GLRLM\_RunPercentage', 'waveletHHL\_GLRLM\_ShortRunEmphasis', 'waveletHHL\_GLRLM\_ShortRunHighGrayLevelEmphasis', 'waveletHHL\_GLRLM\_ShortRunLowGrayLevelEmphasis', 'waveletHHL\_firstOrder\_CoefficientVariation', 'waveletHHL\_firstOrder\_Energy', 'waveletHHL\_firstOrder\_Entropy', 'waveletHHL\_firstOrder\_InterquartileRange', 'waveletHHL\_firstOrder\_Kurtosis', 'waveletHHL\_firstOrder\_Maximum', 'waveletHHL\_firstOrder\_Mean', 'waveletHHL\_firstOrder\_MeanAbsoluteDeviation', 'waveletHHL\_firstOrder\_Median', 'waveletHHL\_firstOrder\_MedianAbsoluteDeviation', 'waveletHHL\_firstOrder\_Minimum', 'waveletHHL\_firstOrder\_Percentile10', 'waveletHHL\_firstOrder\_Percentile90', 'waveletHHL\_firstOrder\_QuartileCoefficientDispersion', 'waveletHHL\_firstOrder\_Range', 'waveletHHL\_firstOrder\_RobustMeanAbsoluteDeviation', 'waveletHHL\_firstOrder\_RootMeanSquare', 'waveletHHL\_firstOrder\_Skewness', 'waveletHHL\_firstOrder\_StandardDeviation', 'waveletHHL\_firstOrder\_Uniformity', 'waveletHHL\_firstOrder\_Variance', 'waveletHLH\_GLCM\_Autocorrelation', 'waveletHLH\_GLCM\_ClusterProminence', 'waveletHLH\_GLCM\_ClusterShade', 'waveletHLH\_GLCM\_ClusterTendency', 'waveletHLH\_GLCM\_Contrast', 'waveletHLH\_GLCM\_Correlation', 'waveletHLH\_GLCM\_DifferenceEntropy', 'waveletHLH\_GLCM\_DifferenceVariance', 'waveletHLH\_GLCM\_Dissimilarity', 'waveletHLH\_GLCM\_Energy', 'waveletHLH\_GLCM\_Entropy', 'waveletHLH\_GLCM\_Homogeneity1', 'waveletHLH\_GLCM\_Homogeneity2', 'waveletHLH\_GLCM\_IDMN', 'waveletHLH\_GLCM\_IDN', 'waveletHLH\_GLCM\_IMC1', 'waveletHLH\_GLCM\_IMC2', 'waveletHLH\_GLCM\_InverseVariance', 'waveletHLH\_GLCM\_MaximumProbability', 'waveletHLH\_GLCM\_SumAverage', 'waveletHLH\_GLCM\_SumEntropy', 'waveletHLH\_GLCM\_SumVariance', 'waveletHLH\_GLCM\_Variance', 'waveletHLH\_GLRLM\_GrayLevelNonUniformity', 'waveletHLH\_GLRLM\_HighGrayLevelRunEmphasis', 'waveletHLH\_GLRLM\_LongRunEmphasis', 'waveletHLH\_GLRLM\_LongRunHighGrayLevelEmphasis', 'waveletHLH\_GLRLM\_LongRunLowGrayLevelEmphasis', 'waveletHLH\_GLRLM\_LowGrayLevelRunEmphasis', 'waveletHLH\_GLRLM\_RunLengthNonUniformity', 'waveletHLH\_GLRLM\_RunPercentage', 'waveletHLH\_GLRLM\_ShortRunEmphasis', 'waveletHLH\_GLRLM\_ShortRunHighGrayLevelEmphasis', 'waveletHLH\_GLRLM\_ShortRunLowGrayLevelEmphasis', 'waveletHLH\_firstOrder\_CoefficientVariation', 'waveletHLH\_firstOrder\_Energy', 'waveletHLH\_firstOrder\_Entropy', 'waveletHLH\_firstOrder\_InterquartileRange', 'waveletHLH\_firstOrder\_Kurtosis', 'waveletHLH\_firstOrder\_Maximum', 'waveletHLH\_firstOrder\_Mean', 'waveletHLH\_firstOrder\_MeanAbsoluteDeviation', 'waveletHLH\_firstOrder\_Median', 'waveletHLH\_firstOrder\_MedianAbsoluteDeviation', 'waveletHLH\_firstOrder\_Minimum', 'waveletHLH\_firstOrder\_Percentile10', 'waveletHLH\_firstOrder\_Percentile90', 'waveletHLH\_firstOrder\_QuartileCoefficientDispersion', 'waveletHLH\_firstOrder\_Range', 'waveletHLH\_firstOrder\_RobustMeanAbsoluteDeviation', 'waveletHLH\_firstOrder\_RootMeanSquare', 'waveletHLH\_firstOrder\_Skewness', 'waveletHLH\_firstOrder\_StandardDeviation', 'waveletHLH\_firstOrder\_Uniformity', 'waveletHLH\_firstOrder\_Variance', 'waveletHLL\_GLCM\_Autocorrelation', 'waveletHLL\_GLCM\_ClusterProminence', 'waveletHLL\_GLCM\_ClusterShade', 'waveletHLL\_GLCM\_ClusterTendency', 'waveletHLL\_GLCM\_Contrast', 'waveletHLL\_GLCM\_Correlation', 'waveletHLL\_GLCM\_DifferenceEntropy', 'waveletHLL\_GLCM\_DifferenceVariance', 'waveletHLL\_GLCM\_Dissimilarity', 'waveletHLL\_GLCM\_Energy', 'waveletHLL\_GLCM\_Entropy', 'waveletHLL\_GLCM\_Homogeneity1', 'waveletHLL\_GLCM\_Homogeneity2', 'waveletHLL\_GLCM\_IDMN', 'waveletHLL\_GLCM\_IDN', 'waveletHLL\_GLCM\_IMC1', 'waveletHLL\_GLCM\_IMC2', 'waveletHLL\_GLCM\_InverseVariance', 'waveletHLL\_GLCM\_MaximumProbability', 'waveletHLL\_GLCM\_SumAverage', 'waveletHLL\_GLCM\_SumEntropy', 'waveletHLL\_GLCM\_SumVariance', 'waveletHLL\_GLCM\_Variance', 'waveletHLL\_GLRLM\_GrayLevelNonUniformity', 'waveletHLL\_GLRLM\_HighGrayLevelRunEmphasis', 'waveletHLL\_GLRLM\_LongRunEmphasis', 'waveletHLL\_GLRLM\_LongRunHighGrayLevelEmphasis', 'waveletHLL\_GLRLM\_LongRunLowGrayLevelEmphasis', 'waveletHLL\_GLRLM\_LowGrayLevelRunEmphasis', 'waveletHLL\_GLRLM\_RunLengthNonUniformity', 'waveletHLL\_GLRLM\_RunPercentage', 'waveletHLL\_GLRLM\_ShortRunEmphasis', 'waveletHLL\_GLRLM\_ShortRunHighGrayLevelEmphasis', 'waveletHLL\_GLRLM\_ShortRunLowGrayLevelEmphasis', 'waveletHLL\_firstOrder\_CoefficientVariation', 'waveletHLL\_firstOrder\_Energy', 'waveletHLL\_firstOrder\_Entropy', 'waveletHLL\_firstOrder\_InterquartileRange', 'waveletHLL\_firstOrder\_Kurtosis', 'waveletHLL\_firstOrder\_Maximum', 'waveletHLL\_firstOrder\_Mean', 'waveletHLL\_firstOrder\_MeanAbsoluteDeviation', 'waveletHLL\_firstOrder\_Median', 'waveletHLL\_firstOrder\_MedianAbsoluteDeviation', 'waveletHLL\_firstOrder\_Minimum', 'waveletHLL\_firstOrder\_Percentile10', 'waveletHLL\_firstOrder\_Percentile90', 'waveletHLL\_firstOrder\_QuartileCoefficientDispersion', 'waveletHLL\_firstOrder\_Range', 'waveletHLL\_firstOrder\_RobustMeanAbsoluteDeviation', 'waveletHLL\_firstOrder\_RootMeanSquare', 'waveletHLL\_firstOrder\_Skewness', 'waveletHLL\_firstOrder\_StandardDeviation', 'waveletHLL\_firstOrder\_Uniformity', 'waveletHLL\_firstOrder\_Variance', 'waveletLHH\_GLCM\_Autocorrelation', 'waveletLHH\_GLCM\_ClusterProminence', 'waveletLHH\_GLCM\_ClusterShade', 'waveletLHH\_GLCM\_ClusterTendency', 'waveletLHH\_GLCM\_Contrast', 'waveletLHH\_GLCM\_Correlation', 'waveletLHH\_GLCM\_DifferenceEntropy', 'waveletLHH\_GLCM\_DifferenceVariance', 'waveletLHH\_GLCM\_Dissimilarity', 'waveletLHH\_GLCM\_Energy', 'waveletLHH\_GLCM\_Entropy', 'waveletLHH\_GLCM\_Homogeneity1', 'waveletLHH\_GLCM\_Homogeneity2', 'waveletLHH\_GLCM\_IDMN', 'waveletLHH\_GLCM\_IDN', 'waveletLHH\_GLCM\_IMC1', 'waveletLHH\_GLCM\_IMC2', 'waveletLHH\_GLCM\_InverseVariance', 'waveletLHH\_GLCM\_MaximumProbability', 'waveletLHH\_GLCM\_SumAverage', 'waveletLHH\_GLCM\_SumEntropy', 'waveletLHH\_GLCM\_SumVariance', 'waveletLHH\_GLCM\_Variance', 'waveletLHH\_GLRLM\_GrayLevelNonUniformity', 'waveletLHH\_GLRLM\_HighGrayLevelRunEmphasis', 'waveletLHH\_GLRLM\_LongRunEmphasis', 'waveletLHH\_GLRLM\_LongRunHighGrayLevelEmphasis', 'waveletLHH\_GLRLM\_LongRunLowGrayLevelEmphasis', 'waveletLHH\_GLRLM\_LowGrayLevelRunEmphasis', 'waveletLHH\_GLRLM\_RunLengthNonUniformity', 'waveletLHH\_GLRLM\_RunPercentage', 'waveletLHH\_GLRLM\_ShortRunEmphasis', 'waveletLHH\_GLRLM\_ShortRunHighGrayLevelEmphasis', 'waveletLHH\_GLRLM\_ShortRunLowGrayLevelEmphasis', 'waveletLHH\_firstOrder\_CoefficientVariation', 'waveletLHH\_firstOrder\_Energy', 'waveletLHH\_firstOrder\_Entropy', 'waveletLHH\_firstOrder\_InterquartileRange', 'waveletLHH\_firstOrder\_Kurtosis', 'waveletLHH\_firstOrder\_Maximum', 'waveletLHH\_firstOrder\_Mean', 'waveletLHH\_firstOrder\_MeanAbsoluteDeviation', 'waveletLHH\_firstOrder\_Median', 'waveletLHH\_firstOrder\_MedianAbsoluteDeviation', 'waveletLHH\_firstOrder\_Minimum', 'waveletLHH\_firstOrder\_Percentile10', 'waveletLHH\_firstOrder\_Percentile90', 'waveletLHH\_firstOrder\_QuartileCoefficientDispersion', 'waveletLHH\_firstOrder\_Range', 'waveletLHH\_firstOrder\_RobustMeanAbsoluteDeviation', 'waveletLHH\_firstOrder\_RootMeanSquare', 'waveletLHH\_firstOrder\_Skewness', 'waveletLHH\_firstOrder\_StandardDeviation', 'waveletLHH\_firstOrder\_Uniformity', 'waveletLHH\_firstOrder\_Variance', 'waveletLHL\_GLCM\_Autocorrelation', 'waveletLHL\_GLCM\_ClusterProminence', 'waveletLHL\_GLCM\_ClusterShade', 'waveletLHL\_GLCM\_ClusterTendency', 'waveletLHL\_GLCM\_Contrast', 'waveletLHL\_GLCM\_Correlation', 'waveletLHL\_GLCM\_DifferenceEntropy', 'waveletLHL\_GLCM\_DifferenceVariance', 'waveletLHL\_GLCM\_Dissimilarity', 'waveletLHL\_GLCM\_Energy', 'waveletLHL\_GLCM\_Entropy', 'waveletLHL\_GLCM\_Homogeneity1', 'waveletLHL\_GLCM\_Homogeneity2', 'waveletLHL\_GLCM\_IDMN', 'waveletLHL\_GLCM\_IDN', 'waveletLHL\_GLCM\_IMC1', 'waveletLHL\_GLCM\_IMC2', 'waveletLHL\_GLCM\_InverseVariance', 'waveletLHL\_GLCM\_MaximumProbability', 'waveletLHL\_GLCM\_SumAverage', 'waveletLHL\_GLCM\_SumEntropy', 'waveletLHL\_GLCM\_SumVariance', 'waveletLHL\_GLCM\_Variance', 'waveletLHL\_GLRLM\_GrayLevelNonUniformity', 'waveletLHL\_GLRLM\_HighGrayLevelRunEmphasis', 'waveletLHL\_GLRLM\_LongRunEmphasis', 'waveletLHL\_GLRLM\_LongRunHighGrayLevelEmphasis', 'waveletLHL\_GLRLM\_LongRunLowGrayLevelEmphasis', 'waveletLHL\_GLRLM\_LowGrayLevelRunEmphasis', 'waveletLHL\_GLRLM\_RunLengthNonUniformity', 'waveletLHL\_GLRLM\_RunPercentage', 'waveletLHL\_GLRLM\_ShortRunEmphasis', 'waveletLHL\_GLRLM\_ShortRunHighGrayLevelEmphasis', 'waveletLHL\_GLRLM\_ShortRunLowGrayLevelEmphasis', 'waveletLHL\_firstOrder\_CoefficientVariation', 'waveletLHL\_firstOrder\_Energy', 'waveletLHL\_firstOrder\_Entropy', 'waveletLHL\_firstOrder\_InterquartileRange', 'waveletLHL\_firstOrder\_Kurtosis', 'waveletLHL\_firstOrder\_Maximum', 'waveletLHL\_firstOrder\_Mean', 'waveletLHL\_firstOrder\_MeanAbsoluteDeviation', 'waveletLHL\_firstOrder\_Median', 'waveletLHL\_firstOrder\_MedianAbsoluteDeviation', 'waveletLHL\_firstOrder\_Minimum', 'waveletLHL\_firstOrder\_Percentile10', 'waveletLHL\_firstOrder\_Percentile90', 'waveletLHL\_firstOrder\_QuartileCoefficientDispersion', 'waveletLHL\_firstOrder\_Range', 'waveletLHL\_firstOrder\_RobustMeanAbsoluteDeviation', 'waveletLHL\_firstOrder\_RootMeanSquare', 'waveletLHL\_firstOrder\_Skewness', 'waveletLHL\_firstOrder\_StandardDeviation', 'waveletLHL\_firstOrder\_Uniformity', 'waveletLHL\_firstOrder\_Variance', 'waveletLLH\_GLCM\_Autocorrelation', 'waveletLLH\_GLCM\_ClusterProminence', 'waveletLLH\_GLCM\_ClusterShade', 'waveletLLH\_GLCM\_ClusterTendency', 'waveletLLH\_GLCM\_Contrast', 'waveletLLH\_GLCM\_Correlation', 'waveletLLH\_GLCM\_DifferenceEntropy', 'waveletLLH\_GLCM\_DifferenceVariance', 'waveletLLH\_GLCM\_Dissimilarity', 'waveletLLH\_GLCM\_Energy', 'waveletLLH\_GLCM\_Entropy', 'waveletLLH\_GLCM\_Homogeneity1', 'waveletLLH\_GLCM\_Homogeneity2', 'waveletLLH\_GLCM\_IDMN', 'waveletLLH\_GLCM\_IDN', 'waveletLLH\_GLCM\_IMC1', 'waveletLLH\_GLCM\_IMC2', 'waveletLLH\_GLCM\_InverseVariance', 'waveletLLH\_GLCM\_MaximumProbability', 'waveletLLH\_GLCM\_SumAverage', 'waveletLLH\_GLCM\_SumEntropy', 'waveletLLH\_GLCM\_SumVariance', 'waveletLLH\_GLCM\_Variance', 'waveletLLH\_GLRLM\_GrayLevelNonUniformity', 'waveletLLH\_GLRLM\_HighGrayLevelRunEmphasis', 'waveletLLH\_GLRLM\_LongRunEmphasis', 'waveletLLH\_GLRLM\_LongRunHighGrayLevelEmphasis', 'waveletLLH\_GLRLM\_LongRunLowGrayLevelEmphasis', 'waveletLLH\_GLRLM\_LowGrayLevelRunEmphasis', 'waveletLLH\_GLRLM\_RunLengthNonUniformity', 'waveletLLH\_GLRLM\_RunPercentage', 'waveletLLH\_GLRLM\_ShortRunEmphasis', 'waveletLLH\_GLRLM\_ShortRunHighGrayLevelEmphasis', 'waveletLLH\_GLRLM\_ShortRunLowGrayLevelEmphasis', 'waveletLLH\_firstOrder\_CoefficientVariation', 'waveletLLH\_firstOrder\_Energy', 'waveletLLH\_firstOrder\_Entropy', 'waveletLLH\_firstOrder\_InterquartileRange', 'waveletLLH\_firstOrder\_Kurtosis', 'waveletLLH\_firstOrder\_Maximum', 'waveletLLH\_firstOrder\_Mean', 'waveletLLH\_firstOrder\_MeanAbsoluteDeviation', 'waveletLLH\_firstOrder\_Median', 'waveletLLH\_firstOrder\_MedianAbsoluteDeviation', 'waveletLLH\_firstOrder\_Minimum', 'waveletLLH\_firstOrder\_Percentile10', 'waveletLLH\_firstOrder\_Percentile90', 'waveletLLH\_firstOrder\_QuartileCoefficientDispersion', 'waveletLLH\_firstOrder\_Range', 'waveletLLH\_firstOrder\_RobustMeanAbsoluteDeviation', 'waveletLLH\_firstOrder\_RootMeanSquare', 'waveletLLH\_firstOrder\_Skewness', 'waveletLLH\_firstOrder\_StandardDeviation', 'waveletLLH\_firstOrder\_Uniformity', 'waveletLLH\_firstOrder\_Variance', 'waveletLLL\_GLCM\_Autocorrelation', 'waveletLLL\_GLCM\_ClusterProminence', 'waveletLLL\_GLCM\_ClusterShade', 'waveletLLL\_GLCM\_ClusterTendency', 'waveletLLL\_GLCM\_Contrast', 'waveletLLL\_GLCM\_Correlation', 'waveletLLL\_GLCM\_DifferenceEntropy', 'waveletLLL\_GLCM\_DifferenceVariance', 'waveletLLL\_GLCM\_Dissimilarity', 'waveletLLL\_GLCM\_Energy', 'waveletLLL\_GLCM\_Entropy', 'waveletLLL\_GLCM\_Homogeneity1', 'waveletLLL\_GLCM\_Homogeneity2', 'waveletLLL\_GLCM\_IDMN', 'waveletLLL\_GLCM\_IDN', 'waveletLLL\_GLCM\_IMC1', 'waveletLLL\_GLCM\_IMC2', 'waveletLLL\_GLCM\_InverseVariance', 'waveletLLL\_GLCM\_MaximumProbability', 'waveletLLL\_GLCM\_SumAverage', 'waveletLLL\_GLCM\_SumEntropy', 'waveletLLL\_GLCM\_SumVariance', 'waveletLLL\_GLCM\_Variance', 'waveletLLL\_GLRLM\_GrayLevelNonUniformity', 'waveletLLL\_GLRLM\_HighGrayLevelRunEmphasis', 'waveletLLL\_GLRLM\_LongRunEmphasis', 'waveletLLL\_GLRLM\_LongRunHighGrayLevelEmphasis', 'waveletLLL\_GLRLM\_LongRunLowGrayLevelEmphasis', 'waveletLLL\_GLRLM\_LowGrayLevelRunEmphasis', 'waveletLLL\_GLRLM\_RunLengthNonUniformity', 'waveletLLL\_GLRLM\_RunPercentage', 'waveletLLL\_GLRLM\_ShortRunEmphasis', 'waveletLLL\_GLRLM\_ShortRunHighGrayLevelEmphasis', 'waveletLLL\_GLRLM\_ShortRunLowGrayLevelEmphasis', 'waveletLLL\_firstOrder\_CoefficientVariation', 'waveletLLL\_firstOrder\_Energy', 'waveletLLL\_firstOrder\_Entropy', 'waveletLLL\_firstOrder\_InterquartileRange', 'waveletLLL\_firstOrder\_Kurtosis', 'waveletLLL\_firstOrder\_Maximum', 'waveletLLL\_firstOrder\_Mean', 'waveletLLL\_firstOrder\_MeanAbsoluteDeviation', 'waveletLLL\_firstOrder\_Median', 'waveletLLL\_firstOrder\_MedianAbsoluteDeviation', 'waveletLLL\_firstOrder\_Minimum', 'waveletLLL\_firstOrder\_Percentile10', 'waveletLLL\_firstOrder\_Percentile90', 'waveletLLL\_firstOrder\_QuartileCoefficientDispersion', 'waveletLLL\_firstOrder\_Range', 'waveletLLL\_firstOrder\_RobustMeanAbsoluteDeviation', 'waveletLLL\_firstOrder\_RootMeanSquare', 'waveletLLL\_firstOrder\_Skewness', 'waveletLLL\_firstOrder\_StandardDeviation', 'waveletLLL\_firstOrder\_Uniformity', 'waveletLLL\_firstOrder\_Variance']

#### Below is the list of gene features

###### ['11-Mar', 'ABCB5', 'ABCC11', 'AC234582.1', 'ACMSD', 'ACSM1', 'ACSM2A', 'ACSM2B', 'ACTG2', 'ACTL6B', 'ADCYAP1R1', 'ADH1A', 'ADRA2C', 'ADRB1', 'AGR3', 'AGRP', 'AGT', 'AGXT', 'AGXT2', 'AHSG', 'AHSP', 'AICDA', 'AIM2', 'AKR1C4', 'ALB', 'ALDH1A3', 'ALDH1L2', 'ALPL', 'ALPPL2', 'ALX1', 'AMBP', 'AMER2', 'AMHR2', 'AMPD1', 'ANO7', 'ANXA10', 'ANXA13', 'ANXA8', 'AOC1', 'APCS', 'APLP1', 'APOA1', 'APOA2', 'APOB', 'APOBEC2', 'APOC3', 'APOD', 'APOH', 'AQP4', 'AR', 'ARG1', 'ARHGAP36', 'ARPP21', 'ASB11', 'ASCL2', 'ASGR2', 'ASIC4', 'ASRGL1', 'ATCAY', 'ATP13A4', 'ATP1A3', 'ATP6V0A4', 'ATP6V0D2', 'ATP6V1B1', 'AXIN2', 'AZGP1', 'AZU1', 'BAALC', 'BAAT', 'BBOX1', 'BCAN', 'BCHE', 'BEND5', 'BEX1', 'BHLHA15', 'BHMT', 'BHMT2', 'BIRC7', 'BLK', 'BMP6', 'BMPER', 'BNC1', 'BPI', 'BPIFB1', 'BRINP1', 'BRINP2', 'BSND', 'BTK', 'C10orf99', 'C11orf21', 'C14orf105', 'C16orf89', 'C18orf42', 'C1QL1', 'C1QTNF4', 'C1orf186', 'C1orf228', 'C1orf61', 'C2orf40', 'C2orf88', 'C4BPA', 'C7', 'C8B', 'C8G', 'C8orf46', 'CA12', 'CA14', 'CA3', 'CA9', 'CACNG7', 'CADM3', 'CALB2', 'CALML3', 'CALML5', 'CAPN3', 'CAPN8', 'CAPS', 'CARTPT', 'CBFA2T3', 'CCL13', 'CCL18', 'CCL19', 'CCL20', 'CCL24', 'CCL25', 'CCNA1', 'CCR2', 'CCR9', 'CD177', 'CD19', 'CD1A', 'CD1B', 'CD1C', 'CD1E', 'CD22', 'CD3D', 'CD70', 'CD79B', 'CD8B', 'CDC42EP5', 'CDH15', 'CDH16', 'CDH17', 'CDHR5', 'CDK5R2', 'CDKN2A', 'CDO1', 'CDX1', 'CDX2', 'CEACAM3', 'CEACAM5', 'CEACAM6', 'CEACAM7', 'CEACAM8', 'CELF3', 'CES1', 'CFHR1', 'CFTR', 'CH507-42P11.8', 'CHAD', 'CHGA', 'CHGB', 'CHRM1', 'CHRNA2', 'CHRNA3', 'CHST4', 'CILP', 'CITED1', 'CLC', 'CLCNKB', 'CLDN10', 'CLDN16', 'CLDN2', 'CLDN6', 'CLDN8', 'CLEC17A', 'CLECL1', 'CLIC6', 'CLRN3', 'CMTM2', 'CNGB1', 'CNTFR', 'COL11A1', 'COL11A2', 'COL17A1', 'COL26A1', 'COL2A1', 'COMP', 'CORO2B', 'CP', 'CPA4', 'CPB1', 'CPB2', 'CPLX2', 'CPNE4', 'CPNE5', 'CRABP1', 'CRISP3', 'CRP', 'CRTAC1', 'CRYAB', 'CRYGN', 'CST1', 'CTNND2', 'CTSE', 'CTSG', 'CTSV', 'CUBN', 'CUX2', 'CWH43', 'CXCL17', 'CXCL5', 'CXCL6', 'CXCL8', 'CXCR2', 'CYP11A1', 'CYP11B1', 'CYP17A1', 'CYP21A2', 'CYP2E1', 'CYP4B1', 'CYP4F3', 'CYS1', 'CYSLTR1', 'D4S234E', 'DAAM2', 'DACH1', 'DAPL1', 'DBH', 'DCDC2', 'DCSTAMP', 'DCT', 'DCX', 'DDC', 'DEFA3', 'DEFA4', 'DEFB1', 'DERL3', 'DES', 'DGKK', 'DHRS2', 'DKK1', 'DLK1', 'DLX3', 'DLX5', 'DMBT1', 'DMGDH', 'DMRT2', 'DNER', 'DNTT', 'DOK5', 'DPCR1', 'DPEP1', 'DPPA3', 'DPPA4', 'DPPA5', 'DPYS', 'DRD2', 'DRGX', 'DSC3', 'DSG1', 'DSG3', 'DTX1', 'DUSP26', 'DUSP5', 'EDNRB', 'EEF1A2', 'EFHD1', 'EGFL6', 'ELANE', 'ELAVL3', 'ELAVL4', 'EMR3', 'ENPEP', 'ENPP3', 'EPHX3', 'ERP27', 'ESR1', 'EXTL1', 'EYA1', 'F13B', 'F2', 'F7', 'FABP1', 'FABP7', 'FAIM2', 'FAIM3', 'FAM101A', 'FAM107A', 'FAM129C', 'FAM131B', 'FAM163A', 'FAM163B', 'FAM166B', 'FAM19A3', 'FAM19A4', 'FAM3D', 'FAM43B', 'FAM83A', 'FAT2', 'FCGBP', 'FCGR3B', 'FCN1', 'FCRL2', 'FCRL3', 'FCRL5', 'FCRLA', 'FEV', 'FGA', 'FGB', 'FGF18', 'FGF21', 'FGF8', 'FGF9', 'FGFBP1', 'FGG', 'FGL1', 'FLT3', 'FOLH1', 'FOLR1', 'FOSB', 'FOXA1', 'FOXA2', 'FOXA3', 'FOXC1', 'FOXC2', 'FOXD1', 'FOXE1', 'FOXH1', 'FOXI1', 'FOXJ1', 'FOXN1', 'FUT3', 'FXYD2', 'G6PC', 'GABBR2', 'GALNT9', 'GAP43', 'GAPDHS', 'GAPT', 'GATA2', 'GATA3', 'GATA4', 'GATA6', 'GBP6', 'GC', 'GCG', 'GCGR', 'GDAP1L1', 'GDF3', 'GDNF', 'GFAP', 'GFRA1', 'GFRA3', 'GIP', 'GJB1', 'GLT1D1', 'GLYAT', 'GNG3', 'GNG7', 'GPA33', 'GPM6A', 'GPM6B', 'GPR143', 'GPR37L1', 'GPR64', 'GPR97', 'GPRC5B', 'GPRC5D', 'GPX2', 'GREM1', 'GRIA3', 'GSTA1', 'GSTA2', 'GSTA3', 'GUCY2C', 'GYG2', 'GZMK', 'HAND1', 'HAND2', 'HAS1', 'HAS2', 'HBA1', 'HBB', 'HBD', 'HDGFL1', 'HEPACAM', 'HEPACAM2', 'HEPHL1', 'HFE2', 'HIST1H1B', 'HIST1H1D', 'HIST1H1E', 'HIST1H2AB', 'HIST1H2AH', 'HIST1H2AI', 'HIST1H2AJ', 'HIST1H2AL', 'HIST1H2AM', 'HIST1H2BB', 'HIST1H2BE', 'HIST1H2BF', 'HIST1H2BG', 'HIST1H2BH', 'HIST1H2BI', 'HIST1H2BL', 'HIST1H2BO', 'HIST1H3B', 'HIST1H3C', 'HIST1H3D', 'HIST1H3F', 'HIST1H3G', 'HIST1H3I', 'HIST1H3J', 'HIST1H4A', 'HIST1H4B', 'HIST1H4C', 'HIST1H4D', 'HIST1H4E', 'HIST1H4F', 'HIST1H4L', 'HIST2H2AB', 'HIST2H2AC', 'HLA-DOB', 'HMP19', 'HMX1', 'HNF1B', 'HNF4A', 'HOGA1', 'HOXA11', 'HOXA5', 'HOXA9', 'HOXB13', 'HOXB5', 'HOXB6', 'HOXB8', 'HOXB9', 'HOXC10', 'HOXD11', 'HOXD8', 'HP', 'HPX', 'HR', 'HRG', 'HSD11B2', 'HSD3B2', 'HSPA2', 'HSPB2', 'HSPB6', 'IBSP', 'ICAM3', 'IFITM5', 'IGF2', 'IGFBP1', 'IGJ', 'IGLL1', 'IGSF1', 'IL18RAP', 'IL21R', 'IL36G', 'IL36RN', 'INHA', 'INMT', 'INPP5J', 'INS', 'INSC', 'INSM1', 'INSM2', 'IRF4', 'IRX5', 'ISL1', 'ITGA10', 'ITIH1', 'ITIH2', 'ITIH3', 'ITIH5', 'ITLN1', 'ITLN2', 'IVL', 'IYD', 'KCNJ15', 'KCNJ16', 'KCNK15', 'KCNK3', 'KCNQ2', 'KHDC3L', 'KIAA0226L', 'KIAA1324', 'KIF26A', 'KIT', 'KL', 'KLHDC8A', 'KLHL14', 'KLK1', 'KLK3', 'KLK4', 'KLK5', 'KLK6', 'KLK7', 'KLK8', 'KNG1', 'KREMEN2', 'KRT13', 'KRT14', 'KRT15', 'KRT16', 'KRT17', 'KRT20', 'KRT5', 'KRT6A', 'KRT6B', 'KRT6C', 'KRTDAP', 'L1TD1', 'LAMP5', 'LBP', 'LCN12', 'LEFTY1', 'LGALS4', 'LGALS7B', 'LGI3', 'LGR5', 'LHFPL3', 'LIM2', 'LIN28A', 'LMO3', 'LMX1B', 'LPAR3', 'LRMP', 'LRP2', 'LRRC14B', 'LRRC15', 'LRRC26', 'LRRN1', 'LRRN4CL', 'LTB', 'LTF', 'LYG1', 'LYPD1', 'LYPD6B', 'LZTS1', 'MAB21L1', 'MAB21L2', 'MAG', 'MAL', 'MARCO', 'MAT1A', 'MB', 'MECOM', 'MEF2B', 'MEOX1', 'MEP1A', 'MEST', 'MFAP5', 'MGAM', 'MGARP', 'MGP', 'MIA', 'MIOX', 'MITF', 'MIXL1', 'MLANA', 'MLC1', 'MLPH', 'MMP1', 'MMP10', 'MMP12', 'MMP17', 'MMP25', 'MMP3', 'MMP8', 'MMP9', 'MOG', 'MPO', 'MRGPRF', 'MS4A1', 'MS4A3', 'MSLN', 'MSMB', 'MSX1', 'MT3', 'MTRNR2L12', 'MTRNR2L8', 'MUC13', 'MUC15', 'MUC16', 'MUC3A', 'MUC5AC', 'MUC6', 'MXRA8', 'MYBPC1', 'MYCN', 'MYH11', 'MYRF', 'MZB1', 'NANOG', 'NANOGP8', 'NAPSA', 'NAT8', 'NCAM1', 'NCAN', 'NDNF', 'NDP', 'NDUFA4L2', 'NEFH', 'NEFL', 'NEFM', 'NFE2', 'NKX2-1', 'NKX3-1', 'NLRP7', 'NNAT', 'NOX1', 'NPTX2', 'NPW', 'NPY', 'NR0B1', 'NR1H4', 'NR5A1', 'NRSN1', 'NTRK1', 'NTRK2', 'NTS', 'NXPH1', 'OCA2', 'OLFM4', 'OLIG1', 'OLIG2', 'OMD', 'OR51E1', 'OR51E2', 'ORM1', 'ORM2', 'P2RX1', 'P2RX3', 'P2RX5', 'PADI4', 'PANX3', 'PAX1', 'PAX2', 'PAX3', 'PAX5', 'PAX9', 'PCDH18', 'PCSK2', 'PDE8B', 'PDIA2', 'PDZK1', 'PEBP4', 'PEG10', 'PEG3', 'PENK', 'PF4', 'PGC', 'PGLYRP1', 'PGLYRP2', 'PHGR1', 'PHOSPHO1', 'PHOX2A', 'PHOX2B', 'PHYHIPL', 'PIGR', 'PIP', 'PKNOX2', 'PKP1', 'PLA2G1B', 'PLA2G2A', 'PLA2G2D', 'PLAC8', 'PLG', 'PLP1', 'PMEL', 'PMP2', 'PNCK', 'PNMAL2', 'PNMT', 'PNOC', 'POF1B', 'POSTN', 'POU2AF1', 'POU2F2', 'POU3F3', 'POU5F1', 'PPP1R1A', 'PRAC1', 'PRAC2', 'PRAM1', 'PRAME', 'PRDM14', 'PRG4', 'PRLHR', 'PRLR', 'PROC', 'PRPH', 'PRSS1', 'PRSS16', 'PRSS3', 'PRSS35', 'PRSS57', 'PRTN3', 'PSMB11', 'PTCRA', 'PTGDS', 'PTGER3', 'PTH1R', 'PTHLH', 'PTPRN', 'PTPRZ1', 'PVALB', 'RAB37', 'RAB38', 'RAB3B', 'RAB3C', 'RAB44', 'RAG1', 'RAG2', 'RASGRP3', 'RBP4', 'RBP5', 'REEP6', 'REG1A', 'REG4', 'RFX4', 'RGL4', 'RGS13', 'RGS20', 'RHBG', 'RHCG', 'RNASE2', 'RNASE3', 'RNF186', 'RNF43', 'RPRM', 'RRAD', 'RSPO1', 'RTN1', 'RUNDC3A', 'RXRG', 'S100A1', 'S100A12', 'S100A14', 'S100A7', 'S100A8', 'S100A9', 'S100B', 'S100P', 'SAA1', 'SALL1', 'SCARF2', 'SCG2', 'SCG3', 'SCG5', 'SCGB1A1', 'SCGB1D2', 'SCGB2A1', 'SCGB2A2', 'SCGB3A1', 'SCGB3A2', 'SCNN1B', 'SCRG1', 'SCTR', 'SCUBE2', 'SDR16C5', 'SEMA5B', 'SERPINA1', 'SERPINA5', 'SERPINA6', 'SERPINB10', 'SERPINB13', 'SERPINB3', 'SERPINB4', 'SERPINC1', 'SERPIND1', 'SERPINE2', 'SFRP2', 'SFRP5', 'SFTA2', 'SFTA3', 'SFTPA1', 'SFTPA2', 'SFTPB', 'SFTPC', 'SFTPD', 'SH2D1A', 'SHISA2', 'SHISA3', 'SHISA8', 'SIX1', 'SIX2', 'SLAMF7', 'SLC16A12', 'SLC17A1', 'SLC17A3', 'SLC17A7', 'SLC18A1', 'SLC1A2', 'SLC22A11', 'SLC22A2', 'SLC22A31', 'SLC24A2', 'SLC26A3', 'SLC26A4', 'SLC26A7', 'SLC27A6', 'SLC28A1', 'SLC30A2', 'SLC34A2', 'SLC35D3', 'SLC38A3', 'SLC3A1', 'SLC44A4', 'SLC45A2', 'SLC45A3', 'SLC4A1', 'SLC4A4', 'SLC6A13', 'SLC6A17', 'SLC6A3', 'SLC7A3', 'SLC7A4', 'SLC7A8', 'SMTNL2', 'SNAP91', 'SNCA', 'SNCG', 'SNX22', 'SOX1', 'SOX17', 'SOX18', 'SOX8', 'SP7', 'SPDEF', 'SPI1', 'SPIB', 'SPINK1', 'SPINK2', 'SPINK4', 'SPNS3', 'SPOCK3', 'SPON1', 'SPRR1A', 'SPRR1B', 'SPRR2A', 'SPRR2D', 'SPRR2E', 'SPRR3', 'SPX', 'SST', 'ST6GAL2', 'ST6GALNAC1', 'STAP1', 'STAR', 'STC2', 'STEAP2', 'STK32A', 'STMN2', 'STMN4', 'SULT2A1', 'SYP', 'SYT1', 'SYT4', 'TAGLN3', 'TBATA', 'TBX3', 'TCEAL7', 'TCERG1L', 'TCF21', 'TCL1A', 'TCN1', 'TF', 'TFF1', 'TFF2', 'TFF3', 'TFPI2', 'TFR2', 'TG', 'TGM1', 'TH', 'TIMP4', 'TLR10', 'TLX2', 'TM4SF4', 'TMEFF2', 'TMEM100', 'TMEM119', 'TMEM151A', 'TMEM200C', 'TMEM213', 'TMEM238', 'TMEM252', 'TMEM27', 'TMEM52B', 'TMEM72', 'TMPRSS11D', 'TNF', 'TNFRSF13B', 'TNFRSF13C', 'TNFRSF17', 'TNNT1', 'TNR', 'TP63', 'TPO', 'TPSAB1', 'TRABD2A', 'TRABD2B', 'TRIB1', 'TRIM15', 'TRIM31', 'TRIM50', 'TRIM63', 'TRIM71', 'TRIML2', 'TRPM1', 'TRPM8', 'TRPS1', 'TRPV6', 'TSHR', 'TSPAN32', 'TSPAN8', 'TTC9B', 'TTR', 'TTYH1', 'TUBB2B', 'TUBB4A', 'TWIST1', 'TXNDC5', 'TYR', 'TYRP1', 'UGT1A9', 'UGT2A3', 'UGT2B4', 'UGT2B7', 'UNCX', 'UPK1A', 'UPK1B', 'UPK2', 'UPK3A', 'UPK3B', 'UTF1', 'VAT1L', 'VENTX', 'VIL1', 'VPREB1', 'VRTN', 'VSIG2', 'VSTM1', 'VSTM2A', 'VTCN1', 'VTN', 'VWA5B2', 'WASF3', 'WFDC2', 'WNT10A', 'WT1', 'ZBED2', 'ZBP1', 'ZCCHC12', 'ZFP42', 'ZG16B']

#### Number of imaging features

### 539

#### Number of gene features

### 959

### -------------------------------Number of Samples for Training and Testing---------------------------------

##### No. of samples for training:84

##### No. of samples for test:22

#### --------------------------Model Summary-----------------------

##### Model Type : multiTaskLinearModel

###### Cross Validation Metrics:

###### Parameters: cv=3 scoring=neg\_mean\_squared\_error

###### Cross validation score:1.29570891618

##### Model Parameters:

###### normalize:False

###### warm\_start:False

###### selection:cyclic

###### fit\_intercept:True

###### l1\_ratio:0.5

###### max\_iter:1000

###### random\_state:None

###### tol:0.0001

###### copy\_X:True

###### alpha:1.0

#### ----------------Model evaluation for Train data--------------------

##### Min Square Error for the Model

###### MSE of train\_eval set:0.401702380138

###### No. of features showing LOW 'RMSE/Stdev' (<=1.0): 959

###### All such features with their Low 'RMSE/Stdev' values could be found in output file: train\_eval\_multiTaskLinearModel\_Labels\_with\_Low\_Ratio.csv No. of features showing HIGH 'RMSE/Stdev' (>1.0): 0 All such features with their High 'RMSE/Stdev' values could be found in output file: train\_eval\_multiTaskLinearModel\_Labels\_with\_High\_Ratio.csv Model evaluation for Train data for label features showing Low 'RMSE/Stdev' (<=1.0) Content-type: text/html Content-type: text/html ----------------Model evaluation for Test data-------------------- Min Square Error for the Model MSE of test\_eval set:1.25581005127 No. of features showing LOW 'RMSE/Stdev' (<=1.0): 84 All such features with their Low 'RMSE/Stdev' values could be found in output file: test\_eval\_multiTaskLinearModel\_Labels\_with\_Low\_Ratio.csv No. of features showing HIGH 'RMSE/Stdev' (>1.0): 874 All such features with their High 'RMSE/Stdev' values could be found in output file: test\_eval\_multiTaskLinearModel\_Labels\_with\_High\_Ratio.csv Model evaluation for Test data for label features showing Low 'RMSE/Stdev' (<=1.0) Content-type: text/html Content-type: text/html
