## Supplementary_report_HNSCC_4 for "ImaGene: A web-based software platform for tumor radiogenomic evaluation and reporting"

### Radiogenomics Analysis Report

##### 10/08/2021 00:05:21

#### ----------------------------Model inputs-------------------------------

##### Mode:Train

##### Model:DecisionTree

##### Params:default

##### No. of imaging features provided: 540

##### No. of gene features provided:976

##### SampleID check results: 'The SampleIDs match for imaging and gene features'

###### Parameters: cv=3 scoring=neg\_mean\_squared\_error

###### Cross validation score:1.03855822222

##### Model Parameters:

###### presort:False

###### splitter:best

###### min\_impurity\_decrease:0.0

###### max\_leaf\_nodes:None

###### min\_samples\_leaf:1

###### min\_samples\_split:2

###### min\_weight\_fraction\_leaf:0.0

###### criterion:mse

###### random\_state:None

###### min\_impurity\_split:None

###### max\_features:None

###### max\_depth:None

#### ----------------Model evaluation for Train data--------------------

##### Min Square Error for the Model

###### MSE of train\_eval set:0.0

###### No. of features showing LOW 'RMSE/Stdev' (<=1.0): 54

###### All such features with their Low 'RMSE/Stdev' values could be found in output file: train\_eval\_DecisionTree\_Labels\_with\_Low\_Ratio.csv No. of features showing HIGH 'RMSE/Stdev' (>1.0): 0 All such features with their High 'RMSE/Stdev' values could be found in output file: train\_eval\_DecisionTree\_Labels\_with\_High\_Ratio.csv Model evaluation for Train data for label features showing Low 'RMSE/Stdev' (<=1.0) Content-type: text/html Content-type: text/html ----------------Model evaluation for Test data-------------------- Min Square Error for the Model MSE of test\_eval set:2.09923242643 No. of features showing LOW 'RMSE/Stdev' (<=1.0): 5 All such features with their Low 'RMSE/Stdev' values could be found in output file: test\_eval\_DecisionTree\_Labels\_with\_Low\_Ratio.csv No. of features showing HIGH 'RMSE/Stdev' (>1.0): 49 All such features with their High 'RMSE/Stdev' values could be found in output file: test\_eval\_DecisionTree\_Labels\_with\_High\_Ratio.csv Model evaluation for Test data for label features showing Low 'RMSE/Stdev' (<=1.0) Content-type: text/html Content-type: text/html
