## Supplementary_report_HNSCC_3 for "ImaGene: A web-based software platform for tumor radiogenomic evaluation and reporting"

### Radiogenomics Analysis Report

##### 09/08/2021 23:11:32

#### ----------------------------Model inputs-------------------------------

##### Mode:Train

##### Model:LinearRegression

##### Params:default

##### No. of imaging features provided: 540

##### No. of gene features provided:976

##### SampleID check results: 'The SampleIDs match for imaging and gene features'

##### No. of samples: 106

##### performing Stand\_scaler normalization for imaging features

##### performing Stand\_scaler for gene features

#### --------------------------Multivariate Correlations (pearson based)-----------------------


### ----------------------------------Features with highly significant correlations-------------------------------------

#### Below is the list of imaging features

###### ['GLRLM\_LongRunLowGrayLevelEmphasis', 'GLRLM\_RunLengthNonUniformity', 'GLSZM\_LZHGE', 'GLSZM\_LZLGE', 'globalHistogram\_Mode', 'shapeSize\_Compactness2', 'shapeSize\_SurfaceArea', 'waveletHLH\_firstOrder\_CoefficientVariation', 'waveletLHH\_GLRLM\_LongRunLowGrayLevelEmphasis', 'waveletLLH\_firstOrder\_CoefficientVariation', 'waveletLLH\_firstOrder\_Median', 'waveletLLL\_GLRLM\_LongRunEmphasis']

#### Below is the list of gene features

###### ['ATP6V0D2', 'C18orf42', 'CYP17A1', 'DCSTAMP', 'IGLL1', 'INSC', 'ITLN1', 'LIN28A', 'MAL', 'MMP8', 'MYH11', 'NANOG', 'NPY', 'PRDM14', 'SERPINA1', 'SLC26A7', 'SP7', 'TMEM52B', 'TSHR', 'TYR', 'VTCN1']

#### Number of imaging features

### 12

#### Number of gene features

### 21

### -------------------------------Number of Samples for Training and Testing---------------------------------

##### No. of samples for training:84

##### No. of samples for test:22

#### --------------------------Model Summary-----------------------

##### Model Type : LinearRegression

###### Cross Validation Metrics:

###### Parameters: cv=3 scoring=neg\_mean\_squared\_error

###### Cross validation score:2.16718355799

##### Model Parameters:

###### copy\_X:True

###### normalize:False

###### n\_jobs:None

###### fit\_intercept:True

#### ----------------Model evaluation for Train data--------------------

##### Min Square Error for the Model

###### MSE of train\_eval set:0.302017441903

###### No. of features showing LOW 'RMSE/Stdev' (<=1.0): 21

###### All such features with their Low 'RMSE/Stdev' values could be found in output file: train\_eval\_LinearRegression\_Labels\_with\_Low\_Ratio.csv No. of features showing HIGH 'RMSE/Stdev' (>1.0): 0 All such features with their High 'RMSE/Stdev' values could be found in output file: train\_eval\_LinearRegression\_Labels\_with\_High\_Ratio.csv Model evaluation for Train data for label features showing Low 'RMSE/Stdev' (<=1.0) Content-type: text/html Content-type: text/html ----------------Model evaluation for Test data-------------------- Min Square Error for the Model MSE of test\_eval set:0.572402014754 No. of features showing LOW 'RMSE/Stdev' (<=1.0): 1 All such features with their Low 'RMSE/Stdev' values could be found in output file: test\_eval\_LinearRegression\_Labels\_with\_Low\_Ratio.csv No. of features showing HIGH 'RMSE/Stdev' (>1.0): 20 All such features with their High 'RMSE/Stdev' values could be found in output file: test\_eval\_LinearRegression\_Labels\_with\_High\_Ratio.csv Model evaluation for Test data for label features showing Low 'RMSE/Stdev' (<=1.0) Content-type: text/html Content-type: text/html
