## Supplementary_report_BC_1 for "ImaGene: A web-based software platform for tumor radiogenomic evaluation and reporting"

### Radiogenomics Analysis Report

##### 04/08/2021 03:18:43

#### ----------------------------Model inputs-------------------------------

##### Mode:Train

##### Model:LinearRegression

##### Params:default

##### No. of imaging features provided: 36

##### No. of gene features provided:976

##### SampleID check results: 'The SampleIDs match for imaging and gene features'

##### No. of samples: 89

##### performing Stand\_scaler normalization for imaging features

##### performing Stand\_scaler for gene features

#### --------------------------Multivariate Correlations (pearson based)-----------------------


### ----------------------------------Features with highly significant correlations-------------------------------------

#### Below is the list of imaging features

###### ['Signal\_Enhancement\_Ratio\_(SER)\_(K7)', 'Size-Lesion\_volume\_(S1)', 'Surface\_Area\_(S3)', 'Volume\_of\_most\_enhancing\_voxels\_(S4)']

#### Below is the list of gene features

###### ['ACTG2', 'ANXA10', 'C2orf88', 'CRABP1', 'GFRA3', 'GPRC5D', 'HAND2', 'HAS1', 'IGF2', 'MLC1', 'PRG4', 'S100B', 'SERPINE2', 'SLC22A31', 'SMTNL2', 'TUBB4A', 'VRTN']

#### Number of imaging features

### 4

#### Number of gene features

### 17

### -------------------------------Number of Samples for Training and Testing---------------------------------

##### No. of samples for training:71

##### No. of samples for test:18

#### --------------------------Model Summary-----------------------

##### Model Type : LinearRegression

###### Cross Validation Metrics:

###### Parameters: cv=2 scoring=neg\_mean\_squared\_error

###### Cross validation score:1.58662239033

##### Model Parameters:

###### copy\_X:True

###### normalize:False

###### n\_jobs:None

###### fit\_intercept:True

#### ----------------Model evaluation for Train data--------------------

##### Min Square Error for the Model

###### MSE of train\_eval set:0.251854599778

###### No. of features showing LOW 'RMSE/Stdev' (<=1.0): 17

###### All such features with their Low 'RMSE/Stdev' values could be found in output file: train\_eval\_LinearRegression\_Labels\_with\_Low\_Ratio.csv No. of features showing HIGH 'RMSE/Stdev' (>1.0): 0 All such features with their High 'RMSE/Stdev' values could be found in output file: train\_eval\_LinearRegression\_Labels\_with\_High\_Ratio.csv Model evaluation for Train data for label features showing Low 'RMSE/Stdev' (<=1.0) Content-type: text/html Content-type: text/html ----------------Model evaluation for Test data-------------------- Min Square Error for the Model MSE of test\_eval set:1.04934720124 No. of features showing LOW 'RMSE/Stdev' (<=1.0): 3 All such features with their Low 'RMSE/Stdev' values could be found in output file: test\_eval\_LinearRegression\_Labels\_with\_Low\_Ratio.csv No. of features showing HIGH 'RMSE/Stdev' (>1.0): 14 All such features with their High 'RMSE/Stdev' values could be found in output file: test\_eval\_LinearRegression\_Labels\_with\_High\_Ratio.csv Model evaluation for Test data for label features showing Low 'RMSE/Stdev' (<=1.0) Content-type: text/html Content-type: text/html
